# Altered Neural Processing in Middle Frontal Gyrus and Cerebellum During Temporal Recalibration of Action-Outcome Predictions in Schizophrenia Spectrum Disorders

**DOI:** 10.1101/2025.04.07.647511

**Authors:** Christina V. Schmitter, Benjamin Straube

**Affiliations:** Department of Psychiatry and Psychotherapy, University of Marburg, Rudolf-Bultmann-Strasse 8, 35039 Marburg, Germany

**Keywords:** sensorimotor temporal recalibration, cross-modal temporal recalibration, multisensory processing, sensorimotor adaptation, prediction, forward model, functional magnetic resonance imaging, schizophrenia

## Abstract

A key function of the perceptual system is to predict the (multi)sensory outcomes of actions and recalibrate these predictions in response to changing conditions. In schizophrenia spectrum disorders (SSD), impairments in this ability have been linked to difficulties in self-other distinction. This study investigated the neural correlates of the recalibration of action-outcome predictions to delays, the transfer of this process across sensory modality, and whether patients with SSD exhibit alterations in the underlying neural processes.

Patients and healthy controls (HC) underwent fMRI while exposed to delays between active or passive button presses and auditory outcomes. A delay detection task assessed recalibration effects on auditory perception (unimodal trials) and its transfer to visual perception (cross-modal trials).

In unimodal trials, HC exhibited reduced activation in left middle frontal gyrus (MFG) after recalibration, particularly for active movements, whereas this effect was reversed in SSD. In cross-modal trials, recalibration was linked to increased activation in bilateral cerebellum in HC, especially for active movements, a pattern significantly reduced in SSD.

These findings suggest that unimodal temporal recalibration of action-outcome predictions in HC is reflected in reduced prediction error-related MFG activity, which is significantly reduced in SSD revealing potentially disrupted recalibration processes. Additionally, cerebellar engagement appears crucial for cross-modal transfer of recalibrated action-outcome timings, a process that may be impaired in SSD, leading to severe perceptual disturbances like hallucinations.

## 1 Introduction

Hallucinations and ego-disturbances are core symptoms of schizophrenia spectrum disorders (SSD). These symptoms have been linked to impairments in the ability to predict the sensory consequences of one’s own actions.^1^ Prominent theories of sensorimotor control propose that internal forward models use copies of the motor commands of our actions to generate predictions about the sensory outcomes these action will produce. Sensory input that aligns with these predictions typically induces modulations in perceptual sensitivity and neural activity across multiple brain regions, thereby facilitating the distinction between self-generated and externally produced sensory signals.^2–7^ However, when this predictive mechanism is disrupted, self-generated sensations may be misattributed to an external source, contributing to the characteristic symptoms of SSD.^8–13^

Crucially, forward model predictions must remain adaptable to maintain an accurate distinction between self- and externally generated sensory input, even in dynamic environments. For example, under certain circumstances, the sensory outcome of an action can be delayed. Research in healthy individuals has consistently shown that repeated exposure to additional action-outcome delays gradually shifts the predicted timing of sensory action-outcomes, so that the delayed outcome is perceived as occurring in synchrony with the action (known as the sensorimotor temporal recalibration effect, TRE).^14–20^

Previous research showed that individuals with SSD exhibit deficits in recalibrating the forward model in response to spatially shifted or rotated visual action feedback.^21–26^ However, evidence regarding temporal recalibration of the forward model in response to delayed action-outcomes in SSD remains less clear. A recent study from our lab found that patients did not exhibit a reduced TRE compared to healthy controls (HC).^27^ Nevertheless, dysfunctions in sensorimotor temporal recalibration may be more subtle and therefore not detectable by conventional binary-choice perceptual tasks, such as synchrony judgment or delay detection tasks. Instead, aberrant processes during sensorimotor temporal recalibration may be reflected in altered neural processing, which may not manifest in perceptual tasks but could still contribute to perceptual disturbances in daily life or increase the likelihood of their occurrence.

Candidate brain regions for aberrant neural processing during temporal recalibration in SSD include regions generally involved in sensorimotor control, such as the cerebellum and the supplementary motor area (SMA). The cerebellum, in particular, has been proposed to play a crucial role in the forward model operations of generating and updating predictions about sensory action-outcomes^3,7,28–32^ and previous studies showed that non-invasive brain stimulation of this region can modulate temporal recalibration.^27,33,34^ Given that cerebellar dysfunctions have been linked to various symptoms in SSD, including auditory verbal hallucinations,^35,36^ altered neural processing in this region could also directly impact temporal recalibration abilities in these patients. Moreover, sensorimotor temporal recalibration was associated with neural activity in regions implicated in domain-general error processing, including the anterior cingulate cortex (ACC) as well as the superior and middle frontal gyri (SFG and MFG).^37–40^ This suggests that recalibration is not confined to sensorimotor networks but also engages higher-order cognitive processing systems. Given that abnormalities in frontal regions are well-documented in SSD, including reduced gray matter volume,^41,42^ altered gyrification,^43–45^ and reduced functional connectivity to temporal^46^ and parietal^47^ regions, such alterations may also contribute to impairments in temporal recalibration.

In this study, we investigated the neural correlates of sensorimotor temporal recalibration in patients with SSD and matched HC. During fMRI data acquisition, participants were exposed to either delayed or undelayed tones following button presses that were either actively performed or passively induced. Active movements were expected to induce sensorimotor temporal recalibration through forward model mechanisms, while passive movements served as a control condition to account for recalibration effects driven by adjustments in the expected inter-sensory timing between tactile sensation of the button press and the tone. We then used a delay detection task to assess (1) how this delay exposure influenced auditory perception (unimodal context). Furthermore, since previous research suggested that forward model predictions might operate at a supra-modal level,^16,20,48,49^ we investigated (2) whether adaptation effects transfer to visual perception (cross-modal context). Based on our previous study, we expected to observe a behavioral TRE in both groups, with no overall impairment in patients with SSD.^27^ This effect was anticipated to be more pronounced in active compared to passive conditions, as recalibration processes driven by the forward model should be present only during active movements. At the neural level, we expected that HC would show delay-dependent modulations in brain activation as neural correlates of recalibration. Since delayed action-outcomes should produce less prediction error responses after recalibration to the delay, we expected reduced activation in brain regions responsible for prediction and prediction error generation, including the cerebellum and frontal regions. In patients with SSD, these delay-dependent modulations were expected to be less pronounced, with the extent of reduction correlating with the severity of positive symptoms, such as hallucinations and delusions.

## 2 Materials and methods

### 2.1 Participants

Twenty-four patients with SSD and 19 HC (10 female, mean age = 37.89 years, *SD* = 10.28) matched for age, sex, and education participated in the study (see **Table 1** for details). Two patients had to be excluded (one due to > 50% missing responses, and one due to technical issues during data collection), resulting in a final sample of 22 patients (11 female, mean age = 35.54 years, *SD* = 11.01). SSD patients were diagnosed with an ICD-10 diagnosis of schizophrenia (NF20 = 13), schizoaffective disorder (NF25 = 7), or acute polymorph psychotic disorder (NF23 = 2). All diagnoses were validated with the German version of the Structured Clinical Interview for DSM-4 (SCID-4). HC did not report any history of psychiatric disorders, as validated with a SCID-4 screening, and reported no first-degree relatives with diagnosed SSD. All participants had normal or corrected-to-normal visual acuity, normal hearing, and no history of neurological disorders. Fourteen patients and 18 HC participated in the experiment during fMRI data acquisition. From the remaining participants, behavioral data were collected outside the MRI scanner as they declined to be scanned or because they met exclusion criteria for MRI (e.g., metallic implants). All participants gave written informed consent and were financially reimbursed for their participation. The study was conducted according to the Declaration of Helsinki and was approved by the local ethics commission (Study 06/19) of the medical faculty of University of Marburg, Germany. The study was pre-registered in the German Clinical Trials Register (DRKS-ID: DRKS00025885; https://drks.de; date of registration: July 23, 2021).

**Table 1.**
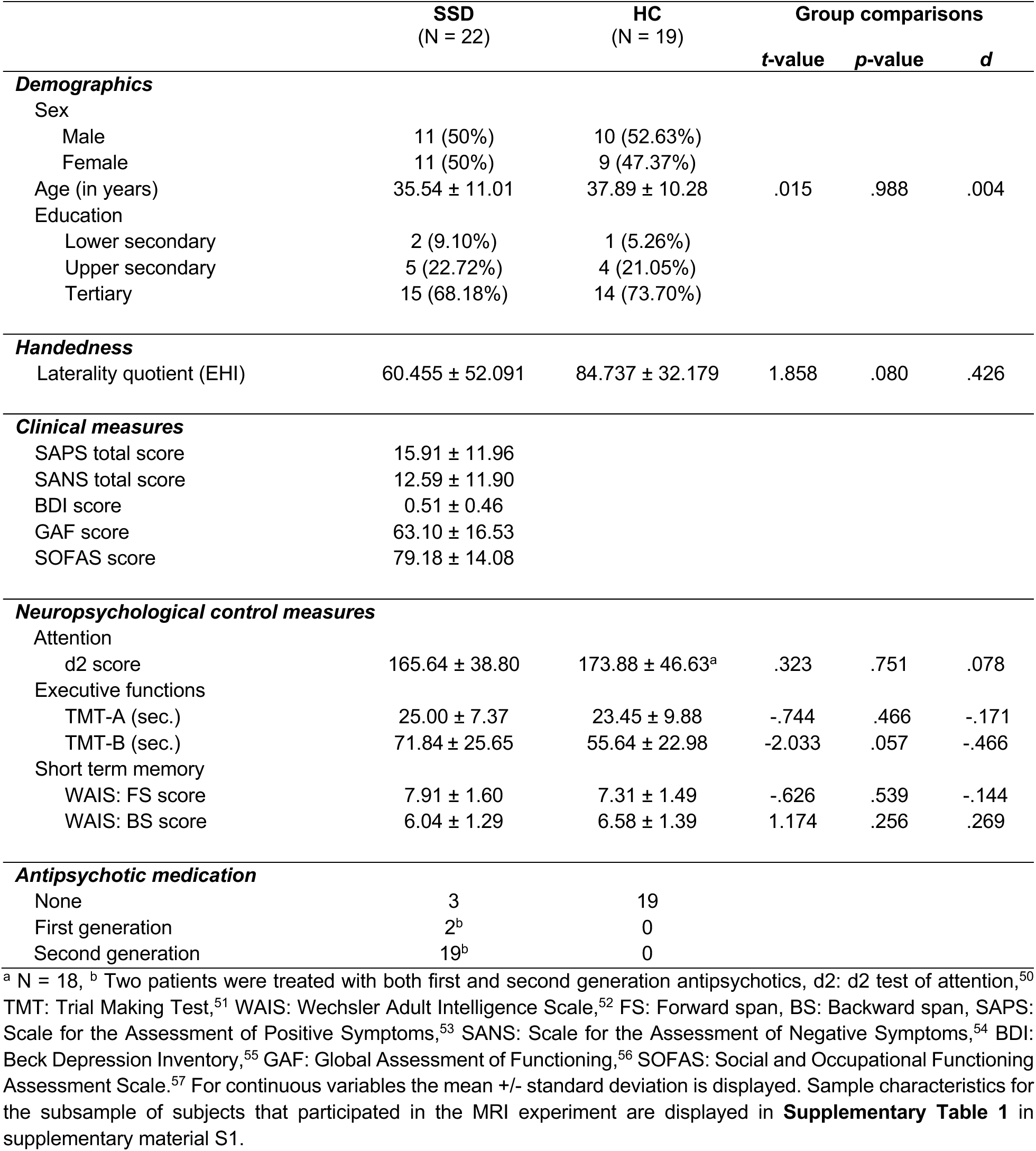
Sample characteristics.

|  | SSD<br>(N = 22) | HC<br>(N = 19) | Group comparisons |  |  |
| --- | --- | --- | --- | --- | --- |
|  |  |  | t-value | p-value | d |
| **Demographics** |  |  |  |  |  |
| Sex |  |  |  |  |  |
| Male | 11 (50%) | 10 (52.63%) |  |  |  |
| Female | 11 (50%) | 9 (47.37%) |  |  |  |
| Age (in years) | 35.54 ± 11.01 | 37.89 ± 10.28 | .015 | .988 | .004 |
| Education |  |  |  |  |  |
| Lower secondary | 2 (9.10%) | 1 (5.26%) |  |  |  |
| Upper secondary | 5 (22.72%) | 4 (21.05%) |  |  |  |
| Tertiary | 15 (68.18%) | 14 (73.70%) |  |  |  |
| **Handedness** |  |  |  |  |  |
| Laterality quotient (EHI) | 60.455 ± 52.091 | 84.737 ± 32.179 | 1.858 | .080 | .426 |
| **Clinical measures** |  |  |  |  |  |
| SAPS total score | 15.91 ± 11.96 |  |  |  |  |
| SANS total score | 12.59 ± 11.90 |  |  |  |  |
| BDI score | 0.51 ± 0.46 |  |  |  |  |
| GAF score | 63.10 ± 16.53 |  |  |  |  |
| SOFAS score | 79.18 ± 14.08 |  |  |  |  |
| **Neuropsychological control measures** |  |  |  |  |  |
| Attention |  |  |  |  |  |
| d2 score | 165.64 ± 38.80 | 173.88 ± 46.63<sup>a</sup> | .323 | .751 | .078 |
| Executive functions |  |  |  |  |  |
| TMT-A (sec.) | 25.00 ± 7.37 | 23.45 ± 9.88 | -.744 | .466 | -.171 |
| TMT-B (sec.) | 71.84 ± 25.65 | 55.64 ± 22.98 | -2.033 | .057 | -.466 |
| Short term memory |  |  |  |  |  |
| WAIS: FS score | 7.91 ± 1.60 | 7.31 ± 1.49 | -.626 | .539 | -.144 |
| WAIS: BS score | 6.04 ± 1.29 | 6.58 ± 1.39 | 1.174 | .256 | .269 |
| **Antipsychotic medication** |  |  |  |  |  |
| None | 3 | 19 |  |  |  |
| First generation | 2<sup>b</sup> | 0 |  |  |  |
| Second generation | 19<sup>b</sup> | 0 |  |  |  |

<sup>a</sup> N = 18, <sup>b</sup> Two patients were treated with both first and second generation antipsychotics, d2: d2 test of attention,<sup>50</sup> TMT: Trial Making Test,<sup>51</sup> WAIS: Wechsler Adult Intelligence Scale,<sup>52</sup> FS: Forward span, BS: Backward span, SAPS: Scale for the Assessment of Positive Symptoms,<sup>53</sup> SANS: Scale for the Assessment of Negative Symptoms,<sup>54</sup> BDI: Beck Depression Inventory,<sup>55</sup> GAF: Global Assessment of Functioning,<sup>56</sup> SOFAS: Social and Occupational Functioning Assessment Scale.<sup>57</sup> For continuous variables the mean +/- standard deviation is displayed. Sample characteristics for the subsample of subjects that participated in the MRI experiment are displayed in **Supplementary Table 1** in supplementary material S1.

### 2.2 Equipment and stimuli

Participants executed button presses of a custom-built, MR-compatible pneumatic passive button device (see **Fig. 1**). During the fMRI experiment, the device was positioned next to their right leg. In active conditions, they pressed the button voluntarily by themselves, whereas in passive conditions, it was depressed by compressed air (max. force 20N). Active and passive movements elicited similar tactile and proprioceptive sensations, allowing the disentanglement of the recalibration of forward model predictions from the recalibration of the timing between different senses. At the end of a button movement (i.e., when it reached the lowest position), visual or auditory stimuli were displayed with or without delay. The visual stimulus was a Gabor patch (1-degree visual angle, spatial frequency: 2 cycles/degree) at the center of a 60Hz monitor located behind the scanner. The auditory stimulus was a sine-wave tone (2000Hz with 2ms rise and fall) presented through headphones. Stimuli were presented for a duration of 33.4ms. A similar experimental setup and the experimental task as described below have been used previously and described in further detail.^39,40^

**Fig. 1.**
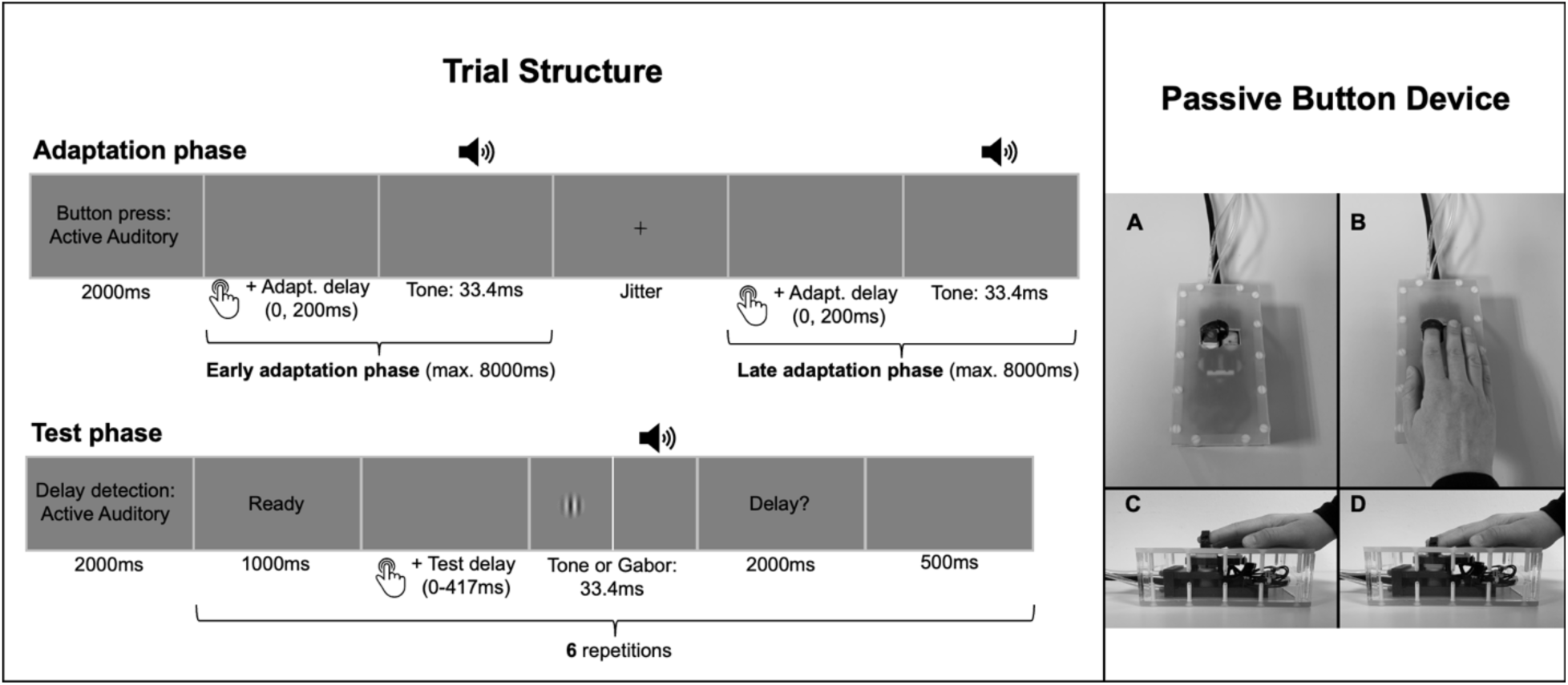
Trial structure and passive button device. Participants completed multiple cycles of adaptation and test phases (see left part of the figure). During adaptation, button presses were either self-initiated or passively induced. Each press triggered a tone, either immediately or with a 200ms delay. Adaptation phases consisted of two segments, separated by a fixation cross of variable duration. In active conditions, participants had 8000ms per part to execute their presses, whereas in passive conditions, nine presses were automatically triggered in each segment. In the subsequent test phase, participants performed a single button press per trial, either actively or passively. The outcome was then presented with one of six possible delays (0–417ms), and they had to indicate whether they perceived a delay. While adaptation always used auditory feedback, the test phase featured either auditory or visual stimuli. Button presses were performed with a custom-built MR-compatible pneumatic passive button device (see right part of the figure). A and B: Participants placed their right index fingers on the button device. The button could be voluntarily pressed by the participants themselves, or it was pulled down passively by compressed air. C: A movement started with the button in the upper position. D: When the button was moved to the lowest position, the stimulus presentation was triggered, either with or without a delay. Data from healthy individuals using the same experimental setup have been published earlier.^39,40^

### 2.3 Experimental design and task description

Participants completed multiple cycles of adaptation and test phases. Each adaptation phase was divided into two segments, separated by the presentation of a fixation cross. In both segments, consecutive button presses were either self-initiated or passively induced (factor *movement type*). Each button press triggered an auditory sensory outcome in the form of a tone. Crucially, during each adaptation phase, the tone was presented consistently either immediately upon button press (undelayed, 0ms delay) or with a delay of 200ms (factor *adaptation delay*). Following each adaptation phase, a test phase evaluated the effect of the previously experienced adaptation delay on perception. Each test phase consisted of six trials, during which the button was pressed once, either actively or passively. The movement type in each test trial always matched that of the preceding adaptation phase. In each trial, the button press triggered the stimulus presentation (visual or auditory, factor *test modality*) with one of six temporal delays (0, 83, 167, 250, 333, or 417ms). Each delay was presented once per test phase in a counterbalanced order. Participants indicated on a button pad, placed on their left leg, whether they perceived a delay between the button press and the corresponding stimulus. The assignment of response options (“delay” or “no delay”) to the response buttons was counterbalanced across participants. The TRE was defined as the difference in delay detection performance between adaptation phases with delayed and undelayed tones. The undelayed tone was considered to align with the natural expectation, while the delayed tones were anticipated to require sensorimotor or inter-sensory temporal recalibration. Lower delay detection rates after exposure to the delayed vs. undelayed tones during adaptation were expected, indicating a shift in the perceived timing of the stimulus toward the adapted delay, thereby reflecting temporal recalibration. The factors *adaptation delay* (0ms vs. 200ms), *movement type* (active vs. passive) and *test modality* (visual vs. auditory) led to an experimental design with eight experimental conditions per *group* (HC vs. SSD).

### 2.4 Procedure

The fMRI experiment consisted of four scanning runs, each comprising 16 cycles of adaptation and test phases. To minimize carryover effects between delayed and undelayed conditions, adaptation delays were blocked within each run. The order of 0ms and 200ms adaptation delay blocks was counterbalanced across participants. Similarly, within each adaptation delay block, active and passive movement conditions were blocked, in counterbalanced order across runs. Each condition appeared in two consecutive cycles of adaptation and test phases per run, totaling eight cycles per condition.

Each adaptation phase began with an instruction screen displayed for 2000ms, indicating the movement type for the upcoming button presses (see **Fig. 1**). Once the instructions disappeared, participants either initiated button presses themselves or experienced passive movements. Each press triggered a tone, either immediately (0ms delay) or with a 200ms delay. In passive conditions, nine button presses were performed in the first part of the adaptation phase, each lasting 500ms with an 800ms interval. This was followed by a fixation cross displayed for a variable duration (1000, 1500, 2000, or 2500ms). Afterward, a second set of nine passive button presses occurred. In active conditions, participants had 8000ms in each part of the adaptation phase to complete their button presses.

Each test phase began with an instruction screen displayed for 2000ms, indicating the movement type and sensory stimulus modality for the upcoming trials. Before each test trial, the cue “Ready” appeared for 1000ms. The trial started as soon as the cue disappeared. In active conditions, participants had 2000ms to execute a single button press. However, participants were instructed to delay their button press by approximately 700ms after the cue disappeared. This was intended to prevent reflexive responses and ensure that the action was genuinely self-initiated.^58^ The onset of passive button movements was jittered (0, 500, or 1000ms). Each button press triggered a visual or auditory outcome with one of six test delay levels. Afterward, the question “Delay?” appeared for 2000ms, prompting participants to indicate via the button pad whether they perceived a delay between the button movement and the outcome. Following a 500ms pause, the “Ready” cue reappeared to start the next trial. The final trial of each test phase was followed by a jittered inter-trial interval (1000, 1500, 2000, or 2500ms) before the adaptation phase of the next cycle started. To ensure task familiarity and correct execution, participants completed a separate training session outside the MRI scanner before the fMRI experiment (see S2 in the supplementary material).

### 2.5 MRI data acquisition

MRI data were collected with a 3 Tesla MR Magnetom TIM Trio scanner (Siemens, Erlangen, Germany) with a 12-channel head-coil at the Department of Psychiatry and Psychotherapy in Marburg. Functional images were obtained parallel to the intercommissural line (anterior commissure – posterior commissure) using a T2*-weighted gradient echo-planar imaging sequence (64 × 64 matrix; repetition time [TR] = 1650ms; echo time [TE] = 25ms; flip angle = 70°; slice thickness = 4.0mm; gap size = 15%; voxel size = 3 × 3 × 4.6mm; field of view [FoV] = 192mm). In each run, 560 volumes of 34 slices each were acquired in descending order covering the whole brain. Anatomical images were obtained using a T1-weighted MPRAGE sequence (256 × 256 matrix; TR = 1900ms; TE = 2.26ms; flip angle = 9°; slice thickness = 1.0mm; gap size = 50%; voxel size = 1 × 1 × 1.5mm; FoV = 256mm). To prevent motion artefacts, subjects’ heads were surrounded by foam pads during data acquisition.

### 2.6 Data analyses

Test trials with incomplete or incorrect movements (i.e., the button failing to reach the lowest position required to trigger stimulus presentation; SSD: 1.95%, HC: 1.44% of all trials) were excluded from behavioral and fMRI analyses. Additionally, trials with missing responses (SSD: 3.74%, HC: 4.47% of all trials) were omitted from the behavioral data analysis. For one HC, an adaptation delay of 150ms was mistakenly used instead of 200ms during data collection. The data of this participant are included in the current analysis.

#### 2.6.1 Analysis of behavioral data

Delay detection performance was assessed based on the proportion of detected delays during test phases, calculated separately for each participant and condition. Psychometric functions, modeled as cumulative Gaussian distributions, were fitted to the data using the Psignifit toolbox (version 4) for Python 3.8.5 (Python Software Foundation, https://www.python.org/). From these functions, delay detection thresholds (the delay detected in 50% of trials), slopes (evaluated at the threshold), and widths (the range between 5% and 95% detection rates) were extracted. Detection thresholds served as an overall measure of performance, with lower values indicating better detection. Slopes and widths reflected the sensitivity to increasing delay levels, capturing the ability to discriminate between different delays. To test for recalibration due to the auditory adaptation delay on auditory outcomes, detection thresholds, slopes, and widths of auditory (unimodal) trials were analyzed using mixed ANOVAs. The model included *group* as a between-participants factor and *adaptation delay* and *movement type* as within-participants factors. The same analysis was conducted for cross-modal (visual) trials to examine modality transfer of recalibration. For significant interactions involving *adaptation delay*, Bonferroni-corrected, two-tailed paired t-tests were conducted to assess TRE differences between groups or movement types. A TRE was defined as an increase in detection thresholds (indicating a shift toward longer delays) or a decrease in slopes and broader widths (reflecting reduced ability to discriminate delay levels) following exposure to delayed versus undelayed tones. Additionally, Bayes factors (BF_inc_l) were computed using Bayesian mixed ANOVAs with default priors. BF_incl_ reflects the ratio of the data’s likelihood under a model including the effect compared to a simpler model without it.^59^ All tests were conducted with JASP (Version 0.19.3).^60^

#### 2.6.2 Analysis of fMRI data

MRI data were preprocessed and analyzed using the Statistical Parametric Mapping toolbox (SPM12; https://www.fil.ion.ucl.ac.uk) in MATLAB (Version 2020b, MathWorks, Sherborn, MA). To account for head motion, functional images were realigned to the mean image of each run. Anatomical images were co-registered to the functional images, segmented, and normalized to the standard Montreal Neurological Institute (MNI) template. Functional images were also normalized to the MNI template, with voxel sizes resampled to 2 × 2 × 2 mm. Finally, spatial smoothing was applied using an 8 mm full-width at half maximum (FWHM) kernel.

A General Linear Model (GLM) was created for each participant. Since we were mainly interested in investigating effects of recalibration on temporal perception, data of the test phases were of primary interest. Therefore, the eight experimental test conditions, defined by *movement type*, *test modality*, and *adaptation delay*, were modeled as separate regressors of interest. Since the study focuses on fMRI activations related to stimulus perception rather than movement execution, test trials were included from stimulus onset to offset, while the remaining trial duration was modeled as a regressor of no interest. Trials without a valid button press were excluded from the analysis. Additionally, data from the adaptation phases were incorporated as regressors of no interest. These were modeled as eight separate regressors based on the factors *movement type*, *adaptation delay*, and *adaptation phase* (early vs. late). Similar to the test phases, adaptation events were included from stimulus onset to offset, with the remaining time between stimuli modeled as a separate regressor of no interest. Adaptation phases without valid button presses in either the early or late segment were excluded from the analysis. The time during the instruction texts, the “Ready” cue, the jitter (fixation cross) in the adaptation phase, and the “Delay?” question were modeled as separate regressors of no interest, as well as the six realignment parameters to account for head motion. A high-pass filter with a 128-second cut-off period was applied to remove low frequencies (<0.0078 Hz) and correct for baseline drifts in the BOLD signal. BOLD responses for all events were modeled using the canonical hemodynamic response function (HRF), with the onset corresponding to each event’s start time. For the single-participant GLMs, T-maps were generated by contrasting each of the eight experimental test conditions against an implicit baseline, which represented the average activation of events not captured by the GLM regressors. For group-level analyses, the resulting contrast estimates from each participant were entered into a flexible factorial design.

To control for multiple comparisons at the cluster level, Monte Carlo simulations^61,62^ were conducted to determine the cluster extent threshold beyond which the false-positive rate remains below alpha = .05 (based on estimated data smoothness of 11mm). After 10,000 simulations, a cluster needed to exceed 99 activated continuous voxels at *p* < .001 uncorrected to achieve correction for multiple comparisons at *p* < .05. Group-level contrast activations were anatomically labeled using the Automated Anatomical Labelling (AAL) toolbox for SPM.^63^

At the group level, hypotheses regarding recalibration-related activations during test phases were tested using T-contrasts. First, we examined auditory (unimodal) trials to assess the effect of recalibration on the perception of the same sensory modality used during adaptation. Next, we analyzed visual (cross-modal) trials to explore the influence of auditory recalibration on visual perception. For both test modalities, we examined contrasts across groups, including the main effect of *adaptation delay* and the interaction between *adaptation delay* and *movement type*. Additionally, we explored group differences, focusing on the two-way interaction between *group* and *adaptation delay*, as well as the three-way interaction involving *group*, *adaptation delay*, and *movement type*. As a sanity check, we also computed the main effect of *movement type* to determine whether stronger motor-related activation was linked to active versus passive movement conditions.

#### 2.6.3 Exploratory correlation analyses of recalibration effects with symptom severity

Since positive symptoms, such as delusions or hallucinations, in SSD have been linked to impaired predictive mechanisms of the forward model, we tested whether patients with more severe positive symptomatology showed a reduced response to the adaptation procedure. To investigate this, we conducted exploratory correlation analyses (uncorrected for multiple comparisons) using the total SAPS score and subscale scores for hallucinations (Scale 1) and delusions (Scale 2). These were correlated with (1) behavioral TREs (separately for active and passive as well as for auditory and visual trials) and (2) neural TREs of the contrast estimates (200ms – 0ms) of each cluster for significant fMRI contrasts involving the *adaptation delay* factor. For main effect contrasts, neural TREs were computed across active and passive conditions. For interaction contrasts that included the factor *movement type*, correlations were computed between symptom scores and neural TREs of active conditions only. This was done to restrict the overall number of correlations and because deficits in the forward model mechanism in patients were expected to specifically affect sensorimotor recalibration as assessed in active conditions. Contrast estimates were extracted as eigenvariates using the VOI function in SPM. Correlations and 90% bootstrapped confidence intervals were computed using the scipy package (version 1.11.1) in Python.

## 3 Results

### 3.1 Behavioral results

#### 3.1.1 Auditory (unimodal) effects

Behavioral results are based on the full sample of 22 SSD and 19 HC. For auditory (unimodal) trials, the mixed ANOVAs revealed significant main effects of *adaptation delay* for the delay detection thresholds [*F*(1, 39) = 40.451, *p* < .001, η^²^_p_ = .509, BF_inc_l = 69.803], with larger thresholds after adaptation to the 200ms (*M* = 280.250, *SE* = 12.602) compared to the 0ms delay (*M* = 253.887, *SE* = 12.547), for the slopes [*F*(1, 39) = 4.242, *p* = .046, η^²^_p_ = .098, BF_incl_ = .453], with flatter slopes after 200ms (*M* = .0049, *SE* < .001) compared to 0ms adaptation (*M* = .0053, *SE* < .001), and for the widths [*F*(1, 39) = 5.899, *p* = .020, η^²^_p_ = .131, BFi_ncl_ = .432], with wider curves after 200ms (*M* = 328.381, *SE* = 16.107) compared to 0ms adaptation (*M* = 303.405, *SE* = 17.612). These effects indicate temporal recalibration, i.e., a shift in temporal perceptual sensitivity towards the delay. For all three measures, no group effects were observed. See **Fig. 2** for a visualization of behavioral effects and S3 in the supplementary material for a detailed summary of all effects and Bayes factors.

**Fig. 2.**
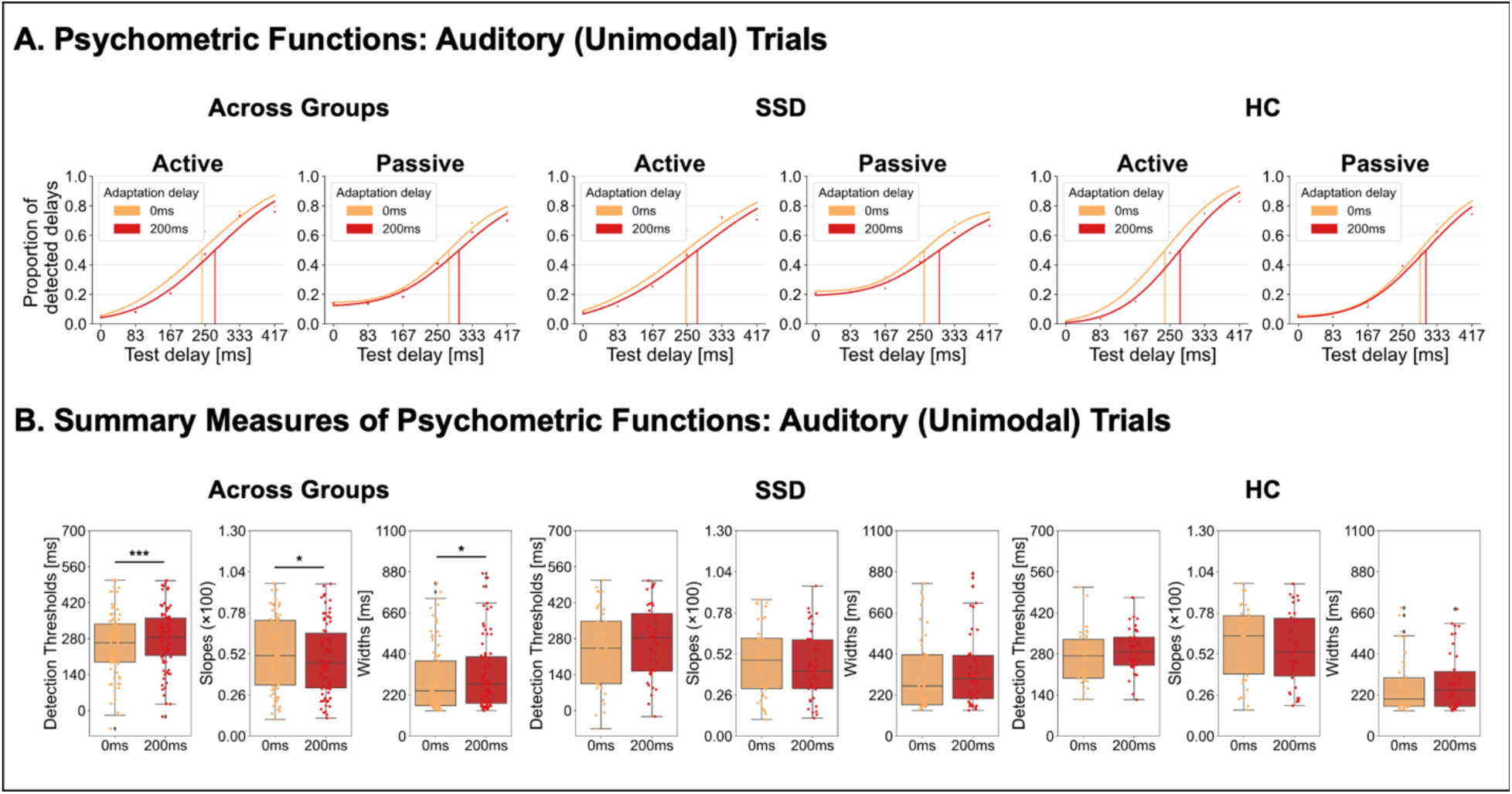
Behavioral results for auditory (unimodal) trials. A. Psychometric functions were fitted to the delay detection data for each experimental condition. B. Across both groups, the TRE manifested as a rightward shift of the functions following exposure to 200ms delayed tones (red) compared to undelayed tones (orange), reflecting increased detection thresholds and thus temporal recalibration. Furthermore, the TRE manifested in flatter slopes and wider curves following delay exposure. For visualization, psychometric functions are shown at the group level, while statistical analyses were based on individually fitted detection rates. * *p* < .05, *** *p* < .001.

#### 3.1.2 Visual (cross-modal) effects

For visual (cross-modal) trials, the mixed ANOVA on delay detection thresholds revealed a significant main effect of *adaptation delay* [*F*(1, 39) = 14.245, *p* < .001, η^²^_p_ = .268, BF_incl_ = 14.131], with larger thresholds after adaptation to the 200ms (*M* = 273.099, *SE* = 13.511) compared to 0ms delay (*M* = 258.189, *SE* = 12.618), indicating a transfer of auditory recalibration effects to vision. Importantly, the significant interaction of *adaptation delay* and *movement type* [*F*(1, 39) = 17.232, *p* < .001, η^²^_p_ = .306, BF_incl_ = 12.428] revealed that this transfer effect was specific to active movements (M_TRE(200-0)_ = 30.313, SE_TRE(200-0)_ = 4.192; *t*(40) = 7.230, *p* < .001, *d* = 1.129, corrected ɑ = 0.025, one-sample, two-sided) but did not occur for passive ones (M_TRE(200-0)_ = .098, SE_TRE(200-0)_ = 6.316; *t*(40) = .016, *p* = .988, *d* = .002, corrected ɑ = 0.025, one-sample, two-sided; active-passive difference: *t*(40) = 4.159, *p* < .001, *d* = .650, paired-samples, two-sided). The ANOVAs on slopes and widths of the curves did not reveal any significant effects with the *adaptation delay* factor. As for the auditory modality, no group effects were observed. See **Fig. 3** for a visualization of behavioral effects for cross-modal trials and S3 in the supplementary material for a detailed summary of all statistics.

**Fig. 3.**
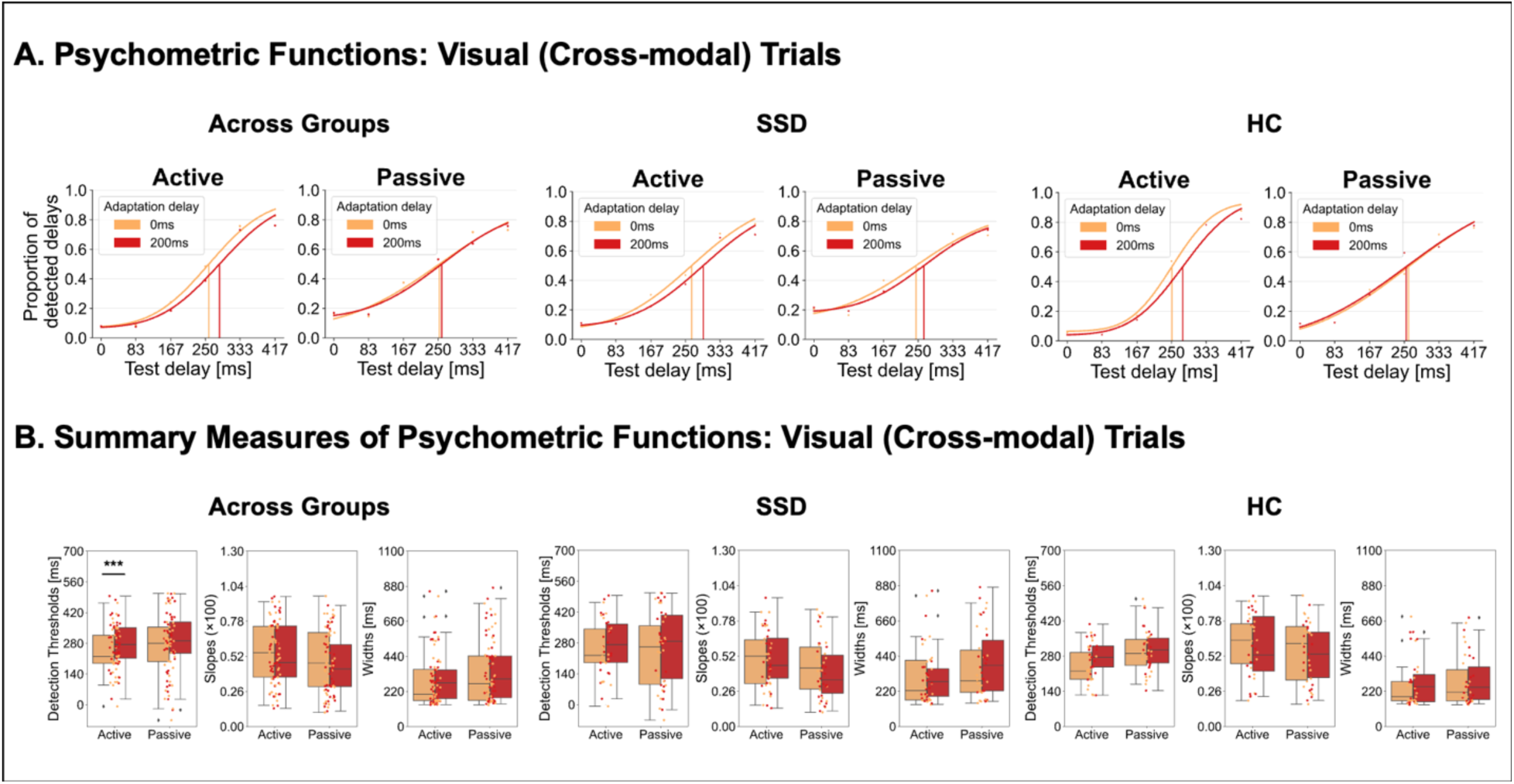
Behavioral results for visual (cross-modal) trials. A. Psychometric functions were fitted to the delay detection data for each experimental condition. B. Across both groups, the TRE manifested as a rightward shift of the functions following exposure to 200ms delayed tones (red) compared to undelayed tones (orange), reflecting increased detection thresholds and thus a transfer of temporal recalibration effects to vision. This transfer effect was specific to active movement conditions. For visualization, psychometric functions are shown at the group level, while statistical analyses were based on individually fitted detection rates. *** *p* < .001.

### 3.2 fMRI results

As a sanity check, we computed the main effect of movement type to determine whether stronger motor-related activation was linked to active versus passive movement conditions. Active movements were associated with stronger activation in left precentral gyrus and the cerebellum (for details see S4 in the supplementary material). This is in line with the idea that forward model predictions based on motor commands were uniquely associated with active conditions in our study.

#### 3.2.1 Commonalities in temporal recalibration across groups

We first examined differences in brain activation across both groups during the auditory (unimodal) delay detection task in test phases, following exposure to delayed versus undelayed tones in the preceding adaptation phases. The 0ms > 200ms contrast identified clusters in left postcentral gyrus, extending into the left superior parietal lobe (SPL), as well as in left SFG, extending to the left SMA and precentral gyrus. These areas exhibited reduced activation during the delay detection task when participants were exposed to delayed tones in the preceding adaptation phase (see **Fig. 4**, left panel and **Table 2**). The reversed contrast (200ms > 0ms) and the interactions of *adaptation delay* (0ms, 200ms) and *movement type* (active, passive) did not reveal significant clusters of activation for auditory trials.

**Fig. 4.**
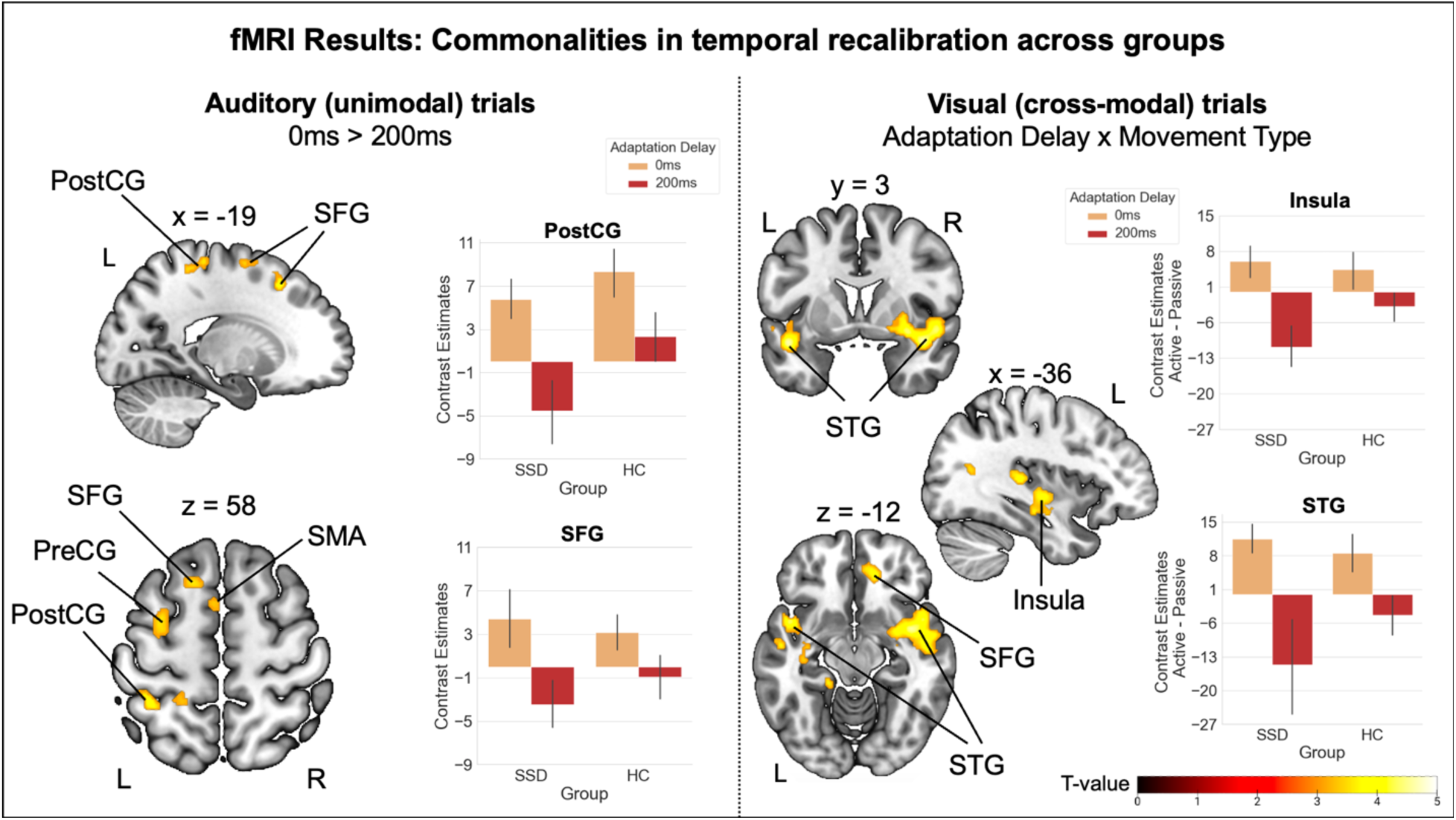
Commonalities in the neural correlates of temporal recalibration across groups. Left panel: Across groups, reduced activation was observed during auditory test phases following exposure to delayed tones during previous adaptation. These effects were found in clusters including the left postcentral gyrus, superior frontal gyrus, and supplementary motor area. **Right panel:** During visual test phases, across groups, reduced activation was observed following exposure to delayed tones in various regions, including superior and middle temporal gyri, right superior frontal gyrus, right putamen and bilateral insula, with this reduction being more pronounced in active than passive conditions. For visualization, contrast estimates (eigenvariates extracted with the VOI function of SPM) are displayed as the difference between active and passive conditions, emphasizing the extent to which the effect is more pronounced in active compared to passive conditions. Error bars show standard errors of the mean. PostCG = postcentral gyrus, PreCG = precentral gyrus, SFG = superior frontal gyrus, SMA = supplementary motor area, STG = superior temporal gyrus. L = left, R = right.

**Table 2.** Group-level contrasts investigating commonalities and differences in temporal recalibration between SSD and HC.

| Cluster peak | Local peaks | Hem. | x | y | z | T-value | no. voxels |
| --- | --- | --- | --- | --- | --- | --- | --- |
| **Commonalities in Auditory Trials: Main Effect of Adaptation Delay (0ms > 200ms)** |  |  |  |  |  |  |  |
| **PostCG** |  | **L** | **-36** | **-44** | **60** | **4.25** | **219** |
|  | PostCG | L | -22 | -30 | 64 | 3.91 |  |
|  | SPG | L | -20 | -42 | 60 | 3.60 |  |
| **SFG** |  | **L** | **-18** | **20** | **50** | **4.22** | **113** |
| **SFG** |  | **L** | **-16** | **4** | **66** | **4.17** | **269** |
|  | SMA | L | -4 | 8 | 60 | 3.74 |  |
|  | PreCG | L | -40 | 0 | 52 | 3.63 |  |
| **Commonalities in Visual Trials: Adaptation Delay x Movement type** |  |  |  |  |  |  |  |
| **TPOsup** |  | **L** | **-46** | **6** | **-16** | **5.09** | **373** |
|  | MTG | L | -52 | -10 | -16 | 4.01 |  |
|  | TPOmid | L | -52 | 10 | -28 | 3.44 |  |
| **PUT** |  | **R** | **32** | **0** | **-8** | **5.02** | **1709** |
|  | INS | R | 42 | -6 | 6 | 4.98 |  |
|  | INS | R | 40 | 2 | -14 | 4.73 |  |
| **HES** |  | **L** | **-30** | **-28** | **6** | **4.97** | **1608** |
|  | tPuM | R | 8 | -24 | 2 | 4.28 |  |
|  | tPuM | L | -4 | -26 | 6 | 4.08 |  |
| **CAL** |  | **R** | **4** | **-64** | **10** | **4.34** | **415** |
|  | CAL | R | 20 | -62 | 10 | 3.38 |  |
| **MTG** |  | **L** | **-56** | **-64** | **16** | **4.19** | **169** |
|  | MOG | L | -48 | -72 | 2 | 3.50 |  |
|  | MTG | L | -36 | -64 | 14 | 3.50 |  |
| **PFCventmed** |  | **R** | **14** | **40** | **-10** | **4.05** | **260** |
|  | ACCsub | L | -6 | 32 | -4 | 3.97 |  |
|  | ACCsub | R | 6 | 32 | -4 | 3.46 |  |
| **INS** |  | **L** | **-36** | **-12** | **-4** | **3.96** | **230** |
|  | HIP | L | -36 | -20 | -14 | 3.74 |  |
|  | INS | L | -42 | -8 | 0 | 3.68 |  |
| **Differences in Auditory Trials: Group x Adaptation Delay x Movement Type** |  |  |  |  |  |  |  |
| **CER VI** |  | **R** | **10** | **-62** | **-24** | **4.48** | **499** |
|  | Vermis VI | . | -4 | -62 | -26 | 3.86 |  |
|  | CER Crus I | L | -16 | -68 | -32 | 3.66 |  |
| **Differences in Visual Trials: Group x Adaptation Delay x Movement Type** |  |  |  |  |  |  |  |
| **MFG** |  | **L** | **-36** | **44** | **18** | **4.06** | **168** |

N<sub>SSD</sub> = 14, N<sub>HC</sub> = 18. Coordinates are listed in MNI space. Significance level: $p < .001$ uncorrected with a minimum cluster extent of 99 voxels ( $p < .05$ Monte Carlo cluster level corrected). PostCG = postcentral gyrus, SPG = superior parietal gyrus, SFG = superior frontal gyrus, SMA = supplementary motor area, PreCG = precentral gyrus, TPOsup = Temporal pole: superior temporal gyrus, MTG = middle temporal gyrus, TPOmid = Temporal pole: middle temporal gyrus, PUT = putamen, INS = insula, HES = Heschl's gyrus, tPuM = Thalamus: pulvinar medial, CAL = Calcarine, MOG = middle occipital gyrus, PFCventmed = Superior frontal gyrus: medial orbital, ACCsub = Anterior cingulate cortex: subgenual, HIP = hippocampus, CER = cerebellum, MFG = middle frontal gyrus, L = left, R = right.

In addition, we investigated the impact of the auditory adaptation procedure on visual perception in cross-modal test phases. Across both groups, no significant main effects of *adaptation delay* were observed. However, the interaction of *adaptation delay* and *movement type* reached significance, with large clusters of activation with peaks in left superior and middle temporal gyri (STG, MTG), right Heschl’s gyrus, right SFG, and right calcarine. Furthermore, significant clusters were identified in subcortical regions, including the right putamen, the bilateral insula, and the left hippocampus (see **Fig. 4**, right panel and **Table 2**). Across both groups, these areas showed reduced activation during the visual delay detection task following prior exposure to delayed tones during adaptation, with this reduction being more pronounced in active compared to passive movement conditions.

#### 3.2.2 Group differences in temporal recalibration

To examine whether delay-related neural activity differed between patients with SSD and HC, we analyzed interaction effects involving *group* and *adaptation delay*. For auditory (unimodal) trials, a significant three-way interaction between *group*, *adaptation delay*, and *movement type* emerged, revealing a cluster of activation in the left MFG. In this region, HC exhibited the previously described pattern of reduced activation during the delay detection task following exposure to delayed tones, with this effect being more pronounced for active than passive movements (see **Fig. 5**, right panel and **Table 2**). In contrast, SSD showed the opposite pattern, displaying increased activation in this cluster after exposure to delayed tones in active movement conditions. No other two- or three-way interaction contrasts with the *group* and *adaptation delay* factors reached significance.

**Fig. 5.**
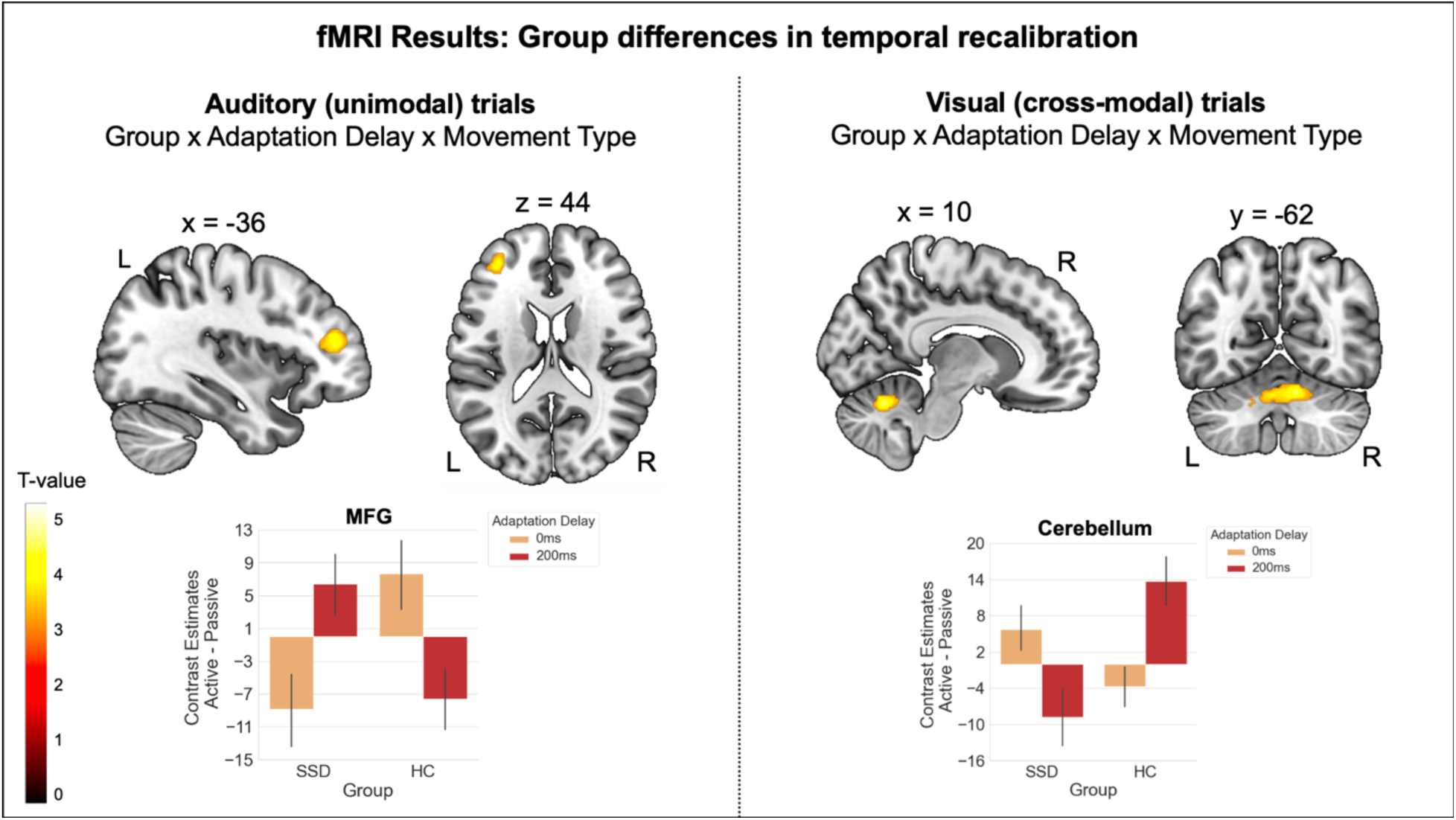
Group differences in the neural correlates of temporal recalibration. Left panel: For auditory (unimodal) trials, group differences in delay-related activation emerged in the left middle frontal gyrus, where HC exhibited reduced activation after delay exposure, particularly in active conditions, whereas SSD showed the opposite pattern. **Right panel:** For visual (cross-modal) trials, group differences in delay-related activation emerged in the cerebellum, where HC exhibited increased activation after delay exposure, particularly in active conditions, while SSD showed the reversed pattern. Error bars show standard errors of the mean. MFG = middle frontal gyrus, L = left, R = right.

As for unimodal trials, we examined group differences in delay-related neural activation between patients with SSD and HC for visual (cross-modal) trials. A significant three-way interaction of *group*, *adaptation delay*, and *movement type* emerged, with a cluster of activation in the right cerebellum (lobule VI), extending into vermis VI and left cerebellar crus I. In this region, HC showed an increase in activation following delay exposure, which was more pronounced in active than in passive conditions (see **Fig. 5**, right panel, and **Table 2**). In contrast, patients with SSD exhibited the opposite pattern. No other two- or three-way interaction contrasts involving the factors *group* and *adaptation delay* reached significance.

### 3.3 Exploratory correlation analyses of recalibration effects with symptom severity

Exploratory correlation analyses were conducted to examine the relationship between symptom severity, as measured by the SAPS total score and subscores for hallucinations and delusions in patients, and both behavioral and neural TREs. For the behavioral TREs, none of the correlations with the symptom scores reached significance for either of the movement types and test modalities (all *p* > .120), indicating that symptom severity was not associated with recalibration, as measured by temporal perceptual sensitivity in the delay detection task. The correlation analyses between symptom severity and the neural TREs (computed as difference in contrast estimates between 200ms and 0ms conditions) revealed significant correlations for the *adaptation delay* x *movement type* interaction in visual (cross-modal) trials. As described in section 3.2.2, this contrast showed reduced activation during the delay detection task after participants were exposed to delayed compared to undelayed tones during adaptation, i.e. negative TRE values, with this effect being particularly pronounced in active conditions. The correlation analyses revealed that patients with a higher SAPS total score and a higher hallucinations subscore were the ones that deviated most strongly from this activation pattern in active conditions, with significant correlations in the cluster with peak activation in putamen (SAPS total score: *r* = .690, *p* = .006, 90% CI [.084, .870]; SAPS hallucination score: *r* = .715, *p* = .004, 90% CI [.386, .865]) and in the cluster with peak activation in Heschl’s gyrus (SAPS total score: *r* = .581, *p* = .029, 90% CI [.075, .831]; SAPS hallucination score: *r* = .611, *p* = .020, 90% CI [.214, .818]). For the other contrasts reported for visual as well as auditory test trials, none of the correlations with symptom severity reached significance. See S5 in the supplementary material for a detailed summary of all correlation results.

## 4 Discussion

This study investigated the neural correlates of sensorimotor temporal recalibration and its cross-modal transfer in patients with SSD and HC. While behavioral results showed comparable recalibration effects in both groups, neural findings revealed key differences in brain activation patterns, particularly in MFG during unimodal recalibration and in cerebellum during cross-modal transfer. These findings provide novel insights into the neural underpinnings of sensorimotor prediction deficits in SSD.

### 4.1 Commonalities in temporal recalibration across groups

At the behavioral level, both groups exhibited significant temporal recalibration effects in unimodal (auditory) conditions, as indicated by increased delay detection thresholds and broader psychometric curves after adaptation to delayed tones. These effects were present for both active and passive conditions, suggesting that they reflect general inter-sensory recalibration mechanisms rather than forward model-specific recalibration.^64,65^ In contrast, cross-modal (visual) recalibration effects were observed only in active conditions, supporting the notion that forward model-based predictions operate at a supra-modal level and influence perception across different sensory modalities. However, it should be noted that heterogeneous findings are reported in the literature regarding action-specific modality transfer and this result should therefore be interpreted with caution.^20,27,39^ Notably, the absence of group differences in behavioral data replicates the previous finding that patients with SSD do not show overt deficits in perceptual temporal recalibration tasks.^27^

At the neural level, in unimodal (auditory) trials, both groups showed reduced activation in postcentral gyrus, SFG, and SMA following adaptation to the delay. These reductions likely reflect a decrease in the amount or amplitude of temporal prediction errors as predictions regarding the timing of the tone shifted towards longer delays and smaller delays are no longer detected. Consequently, after successful recalibration, fewer or smaller prediction errors may have reached sensory regions like the postcentral gyrus to attenuate their activity or frontal areas that are involved in domain-general error processing. It is particularly interesting that activations of this contrast also extended to the SMA, a region implicated in the generation of the efference copy of motor commands that is used by the forward model (potentially located in the cerebellum) for prediction generation.^66^ After the action, prediction errors resulting from the comparison between the forward model prediction and the actual sensory feedback are projected back to the SMA to update the motor plan if necessary.^31,67^ Thus, in our study, after recalibration to the delay, fewer and smaller prediction errors from the cerebellum may have reached the SMA during the delay detection task. This notion is further supported by supplementary parametric analyses of the adaptation phase data, which showed that activation in both the SMA and cerebellum decreased gradually over the course of adaptation (see S6 in the supplementary material). This may reflect a prediction error signal which is stronger in the beginning of adaptation when exposed to the delay but then diminishes as the forward model integrates the delay into its predictions. However, it must be noted that the effect in test phases outlined above was observed across both active and passive conditions, indicating that it may be related to a more domain-general mechanism rather than a specific action-related predictive mechanism.

In cross-modal (visual) trials, a pattern of reduced activation after delay adaptation occurred in both groups, most prominently in STG and Heschl’s gyrus, insula and putamen. Contrary to the pattern observed in unimodal trials, this decrease in activation was particularly pronounced in active conditions. This pattern aligns with the behavioral data, in which a cross-modal transfer of recalibration effects was also observed exclusively in active conditions, suggesting an explicit involvement of action-specific transfer processes of the forward model.^27^ Superior temporal regions, including the STG, are known for their role in cross-modal integration. For instance, activity in this region increases when visual and auditory stimuli are temporally aligned^68,69^ and it is modulated by changes in the audiovisual temporal binding window.^70^ The insula is also considered a key region for multisensory processing.^71,72^ Moreover, it is responsive to prediction errors across various domains, including perceptual errors, where expected sensory input does not match internal predictions.^73,74^ It has been suggested to play a role in integrating body-related information with top-down predictions^75^ and in making the resulting prediction errors accessible to awareness.^76^ Similarly, activity in the putamen has been linked to the processing of prediction errors in action-outcome associations and action-related rewards,^77,78^ as well as in action-independent stimulus associations.^79^ Furthermore, the timing of putamen activation has been connected to delay detection performance, particularly for active as opposed to passive movements in healthy individuals.^80^ Both the insula and the putamen have also been implicated during the pre-movement period, when predictions about upcoming visual feedback are formed.^81^ In our study, the STG as well as the insula and the putamen may play a crucial role in transferring the temporal mapping between movement and tone, learned during the adaptation phase, to visual stimuli. Consequently, these regions show a reduced temporal prediction error response during the delay detection task in the cross-modal context following successful recalibration. Even though the involvement of these regions in cross-modal processes has been identified across domains of action and perception, our results suggest a particularly strong engagement when (potentially supra-modal) action-related temporal adaptation effects are transferred to another modality. Since these effects were observed across both groups, modality transfer processes in these regions appear to remain intact in individuals with SSD.

### 4.2 Altered MFG activity after unimodal recalibration in SSD

Next to recalibration-related neural correlates found across groups, we also assessed group differences in the neural correlates of temporal recalibration. For unimodal trials, group differences emerged in MFG, where HC exhibited reduced activation after adaptation to the delay, particularly in active conditions (evident after subtracting activation from the passive control condition), while SSD patients showed the opposite pattern. Given the role of the MFG in conflict monitoring and predictive control, also during sensorimotor temporal recalibration,^37,39,40^ this suggests that HC successfully recalibrated their forward model predictions, reducing prediction error signals for delays in this region. In contrast, the increased activation in SSD may indicate continued conflict as a cause or consequence of an impaired ability to adequately recalibrate their forward model predictions. This aligns with findings of general frontal deficits in patients which include, e.g., MFG volume reductions^82,83^ and prefrontal-striatal hypoconnectivity.^84,85^ Furthermore, in a recent study, frontal non-invasive brain stimulation via tDCS specifically modulated forward model-related predictive mechanisms in patients,^86^ which further highlights the crucial role of dysfunctional frontal processes in this context. As recalibration deficits did not manifest in the perceptual task in patients, neural activations assessed via fMRI appear to be the more sensitive measure for detecting differences between HC and patients in sensorimotor temporal recalibration.

### 4.3 Impaired cerebellar processing during cross-modal transfer in SSD

While patients and HC shared neural correlates of the modality transfer process in regions like STG, insula and putamen, group differences were observed in the cerebellum. HC exhibited increased activation in this region after adaptation to delays in active conditions, while SSD patients showed the reverse pattern. The cerebellum has frequently been proposed as the key region for instantiating forward models that generate predictions not only about the sensory outcomes of one’s own actions^7,31,87^ but also about action-independent sensory events.^88,89^ It compares these predictions with incoming sensory input^3,32,67^ and updates them in response to repeated prediction errors.^27,34,39,40,90^ Moreover, the cerebellum has often been suggested to generate predictions at a supra-modal level^7,27,34^ (but see^39^) and has also been identified as playing a role in multisensory integration.^72,91,92^ The increased activation of the cerebellum in the visual delay detection task in HC after adaptation to the delayed tone, specifically in active conditions, may therefore indicate greater processing demands in prediction generation after forward model has just been updated.^39^ In our study, this appears to be particularly the case when a transfer or integration effort is required in order to transfer the adapted temporal motor-auditory mapping to the untrained visual modality. Patients with SSD are known to exhibit structural and functional alterations in the cerebellum, such as reduced size and cerebellar blood flow,^35^ as well as decreased structural and resting-state functional connectivity.^36^ Given the cerebellum’s extensive connectivity with various cortical and subcortical regions, such abnormalities may underlie a wide range of symptoms in SSD.^35^ In our study, this appears to affect the cerebellum’s ability to support cross-modal sensorimotor temporal recalibration in patients. As in unimodal trials, alterations in cross-modal recalibration did not manifest in the perceptual task but became evident only in the neural correlates of this process.

### 4.4 Implications for symptomatology in SSD

Exploratory correlation analyses between neural TREs and SAPS scores revealed that patients with higher total and hallucinations SAPS scores exhibited the strongest deviations from the recalibration-related activation pattern observed across both groups in putamen and Heschl’s gyrus. These correlations were exclusively found for the cross-modal (visual) condition. In putamen and Heschl’s gyrus, both groups exhibited reduced activation after delay adaptation in visual trials, particularly in active conditions. This can be explained by a reduction in sensorimotor prediction error responses in these regions, as fewer delays were detected following adaptation. However, in patients with higher symptom severity, particularly hallucinations, greater prediction error signals persisted in these regions. This may reflect either a cause or a consequence of less effective sensorimotor temporal recalibration processes and aligns with previous findings linking alterations in neural processing in these regions to SSD, particularly to the occurrence of hallucinations. For example, patients with SSD have been shown to exhibit reduced preparatory activity in the putamen prior to movement execution compared to HC.^81^ Moreover, numerous studies have reported a reduced volume^93^ and altered connectivity of Heschl’s gyrus,^94^ as well as diminished suppression of self-generated sensory input in patients^95^ – particularly in those experiencing auditory hallucinations. Even though the reported correlations with activity in these regions in our study should be interpreted with caution due to the small sample size, they suggest a direct link between altered neural correlates of forward model recalibration and its modality transfer and the emergence of positive symptoms, particularly hallucinations, in SSD. A reduced flexibility of these predictive processes in recalibrating to dynamically changing environmental conditions, may contribute to difficulties in self-other distinction and, ultimately, to the manifestation of symptoms such as hallucinations.

### 4.5 Limitations

Some limitations of the samples used in the present study should be discussed. First, the majority of patients exhibited only mild to moderate symptom severity, as assessed in the clinical interview. This was largely due to the requirement for a certain level of functioning to participate in the experiment. Consequently, we cannot rule out the possibility that group differences between patients and HC might have emerged not only in the more sensitive brain measures but also in the perceptual task in a sample with higher symptom scores. The reported correlations with symptom severity should therefore also be interpreted with caution and validated in a larger sample. Second, most patients were receiving antipsychotic medication at the time of testing, which may have attenuated dysfunctions in the investigated mechanism, making them more difficult to detect. Importantly, however, there were no correlations between antipsychotic medication and either behavioral or neural markers of temporal recalibration, suggesting that medication itself is unlikely to have biased the reported effects (see S7 in the supplementary material).

### 4.6 Conclusions

Our findings suggest that SSD is associated with specific alterations in the neural correlates of sensorimotor temporal recalibration, despite preserved behavioral performance in the delay detection task. Patients did not show the reduced prediction error-related activity in MFG after recalibration as observed in HC as a potential marker for successful unimodal recalibration. Furthermore, they exhibited a reduced cerebellar engagement required for the transfer of temporal recalibration effects to another sensory modality. These alterations in neural processing during sensorimotor temporal recalibration in SSD may contribute to positive symptoms like hallucinations.

## Acknowledgements

We thank Jens Sommer and Olaf Steinsträter for technical support, Michelle Achenbach, Samentha Dabaree, Trâm Ðỗ, Luca Grolms, Lisa Herberstein, and Sabrina Nasri-Roudsari for assistance in data collection and patient recruitment. Furthermore, we thank Mechthild Wallnig and Rita Werner for assistance in MRI data acquisition.

## Data and code availability statement

The data that support the findings of this study are openly available in Zenodo at: 10.5281/zenodo.15144534

## Ethics approval statement

The study was approved by the local ethics commission (Study 06/19) of the medical faculty of University of Marburg, Germany

## Author contribution statement

BS and CS conceived and designed the experiment and discussed and interpreted the data. BS was responsible for funding acquisition, project administration and supervision. CS was responsible for data acquisition and analysis and wrote the manuscript. Both authors revised and approved the submitted version of the manuscript.

## Supplementary material

### S1. Details on sample characteristics

Out of the total sample of 22 patients and 19 HC, 14 patients and 18 HC completed the experiment during fMRI data acquisition. The remaining participants either declined to be scanned or were unable to undergo MRI due to metallic implants. The patient fMRI sample contained 8 patients with schizophrenia (F20) and 6 patients with schizoaffective disorder (F25). Detailed sample characteristics of the fMRI-subsamples are displayed in **Supplementary Table 1**.

**Supplementary Table 1.**
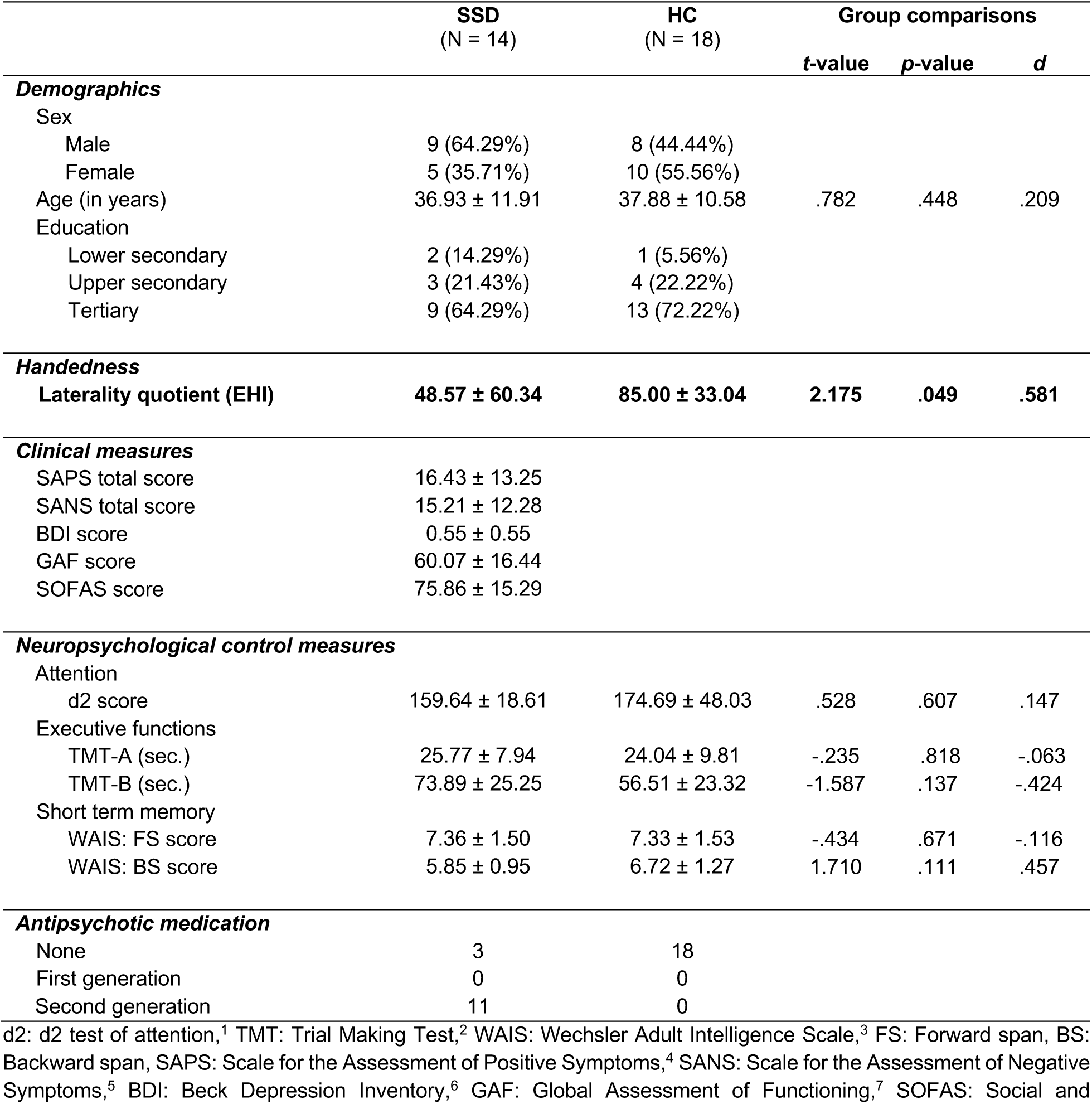
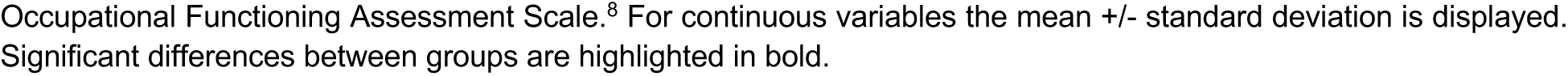
fMRI-Sample characteristics.

### S2. Training Procedure

To ensure correct execution of button presses and familiarize participants with the task, all participants underwent a training session before performing the fMRI experiment. The training procedure for this experimental task has been described in detail previously.^9^ Participants practiced allowing their finger to be moved passively by the button device without applying counter-pressure and learned to execute button presses with precise timing – approx. every 800ms during adaptation phases and lasting around 500ms in both adaptation and test phases. The 500ms duration ensured that stimuli were always presented before the button‘s upward movement, preventing potential interference with delay detection. Each adaptation phase ended automatically after nine button presses per segment. If participants pressed too quickly (faster than 8000ms in total), the inter-phase jitter (if in the first segment) or the instruction display before the test phase (if in the second segment) was extended accordingly. To further familiarize themselves with the task, participants practiced test phases for each condition – once without a delay and once with the maximum delay (417ms). During this training, they received feedback on whether a delay was present. They were instructed to respond as accurately as possible, without time pressure. Finally, they completed a 10-minute training session of the full experiment.

### S3. Detailed summary of behavioral results

To test for recalibration effects after exposure to the auditory adaptation delay on auditory perception (unimodal trials) and its transfer to visual perception (cross-modal trials), detection thresholds, slopes, and widths of the psychometric functions were analyzed using mixed ANOVAs for the respective trials. Furthermore, Bayes factors (BFincl) were calculated for all effects. The results of these analyses are summarized in **Supplementary Tables 1 and 2.**

**Supplementary Table 2.** Results of the ANOVA on detection thresholds, slopes, and widths of the psychometric functions in unimodal trials (auditory test modality).

| Effect | df | F | p | $\eta^2_p$ | $BF_{incl}$ |
| --- | --- | --- | --- | --- | --- |
| **Detection thresholds** |  |  |  |  |  |
| Group | 1 | 0.017 | 0.898 | $4.303 \times 10^{-4}$ | 0.252 |
| Residuals | 39 |  |  |  |  |
| Movement type | 1 | 3.290 | 0.077 | 0.078 | 2.064 |
| Movement type * Group | 1 | 0.186 | 0.669 | 0.005 | 0.191 |
| Residuals | 39 |  |  |  |  |
| **Adaptation delay** | **1** | **40.451** | **< .001** | **0.509** | **69.803** |
| Adaptation delay * Group | 1 | 0.058 | 0.812 | 0.001 | 0.193 |
| Residuals | 39 |  |  |  |  |
| Movement type * Adaptation delay | 1 | 2.398 | 0.130 | 0.058 | 0.685 |
| Movement type * Adaptation delay * Group | 1 | 0.034 | 0.855 | $8.618 \times 10^{-4}$ | 0.023 |
| Residuals | 39 |  |  |  |  |
| **Slopes** |  |  |  |  |  |
| Group | 1 | 0.464 | 0.500 | 0.012 | 0.246 |
| Residuals | 39 |  |  |  |  |
| **Movement type** | **1** | **10.622** | **0.002** | **0.214** | **8.810** |
| Movement type * Group | 1 | 0.842 | 0.364 | 0.021 | 0.290 |
| Residuals | 39 |  |  |  |  |
| **Adaptation delay** | **1** | **4.242** | **0.046** | **0.098** | **0.453** |
| Adaptation delay * Group | 1 | 0.324 | 0.572 | 0.008 | 0.145 |
| Residuals | 39 |  |  |  |  |
| Movement type * Adaptation delay | 1 | 0.456 | 0.503 | 0.012 | 0.314 |
| Movement type * Adaptation delay * Group | 1 | 0.187 | 0.668 | 0.005 | 0.027 |
| Residuals | 39 |  |  |  |  |
| **Widths** |  |  |  |  |  |
| Group | 1 | 0.157 | 0.694 | 0.004 | 0.231 |
| Residuals | 39 |  |  |  |  |
| **Movement type** | **1** | **9.819** | **0.003** | **0.201** | **5.405** |
| Movement type * Group | 1 | 1.010 | 0.321 | 0.025 | 0.341 |
| Residuals | 39 |  |  |  |  |
| **Adaptation delay** | **1** | **5.899** | **0.020** | **0.131** | **0.432** |
| Adaptation delay * Group | 1 | 0.344 | 0.561 | 0.009 | 0.112 |
| Residuals | 39 |  |  |  |  |
| Movement type * Adaptation delay | 1 | 0.021 | 0.886 | $5.332 \times 10^{-4}$ | 0.213 |
| Movement type * Adaptation delay * Group | 1 | 0.579 | 0.451 | 0.015 | 0.019 |
| *Residuals* | 39 |  |  |  |  |

N<sub>SSD</sub> = 22, N<sub>HC</sub> = 19. Significant effects ( $p < .05$ ) are highlighted in bold.

**Supplementary Table 3.**
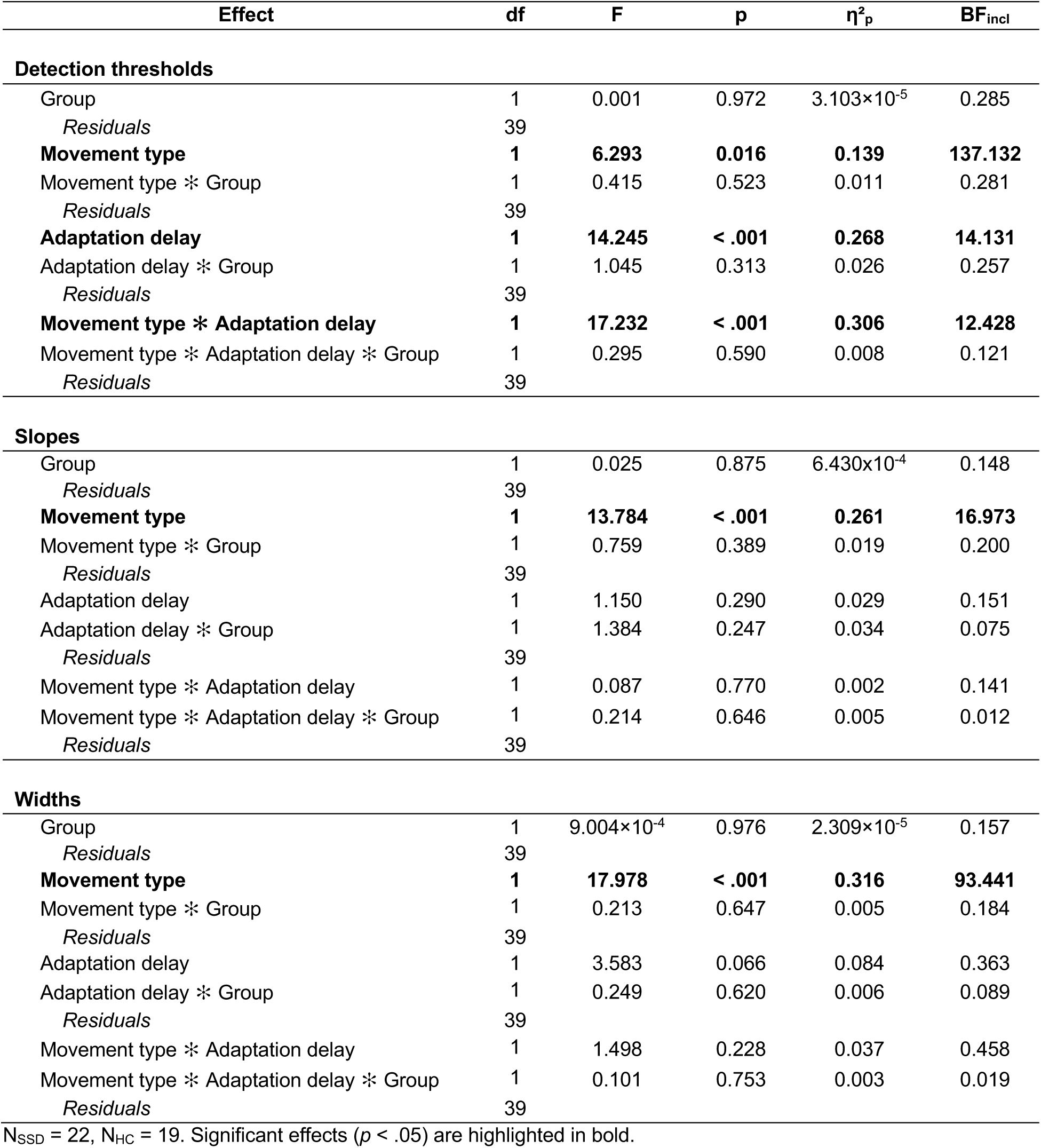
Results of the ANOVA on detection thresholds, slopes, and widths of the psychometric functions in cross-modal trials (visual test modality).

### S4. Main effect of Movement Type (Active > Passive)

To determine whether motor-related processes were specific to active movement conditions, we compared brain activation between active and passive conditions (Active > Passive) in test phases. As expected, active conditions elicited stronger activation in motor-related regions, particularly in the left precentral gyrus and the cerebellum (see **Supplementary Fig. 1** and **Supplementary Table 4**). This suggests that motor-related predictive processes that were of primary interest in our study should be specific to active conditions.

**Supplementary Fig. 1.**
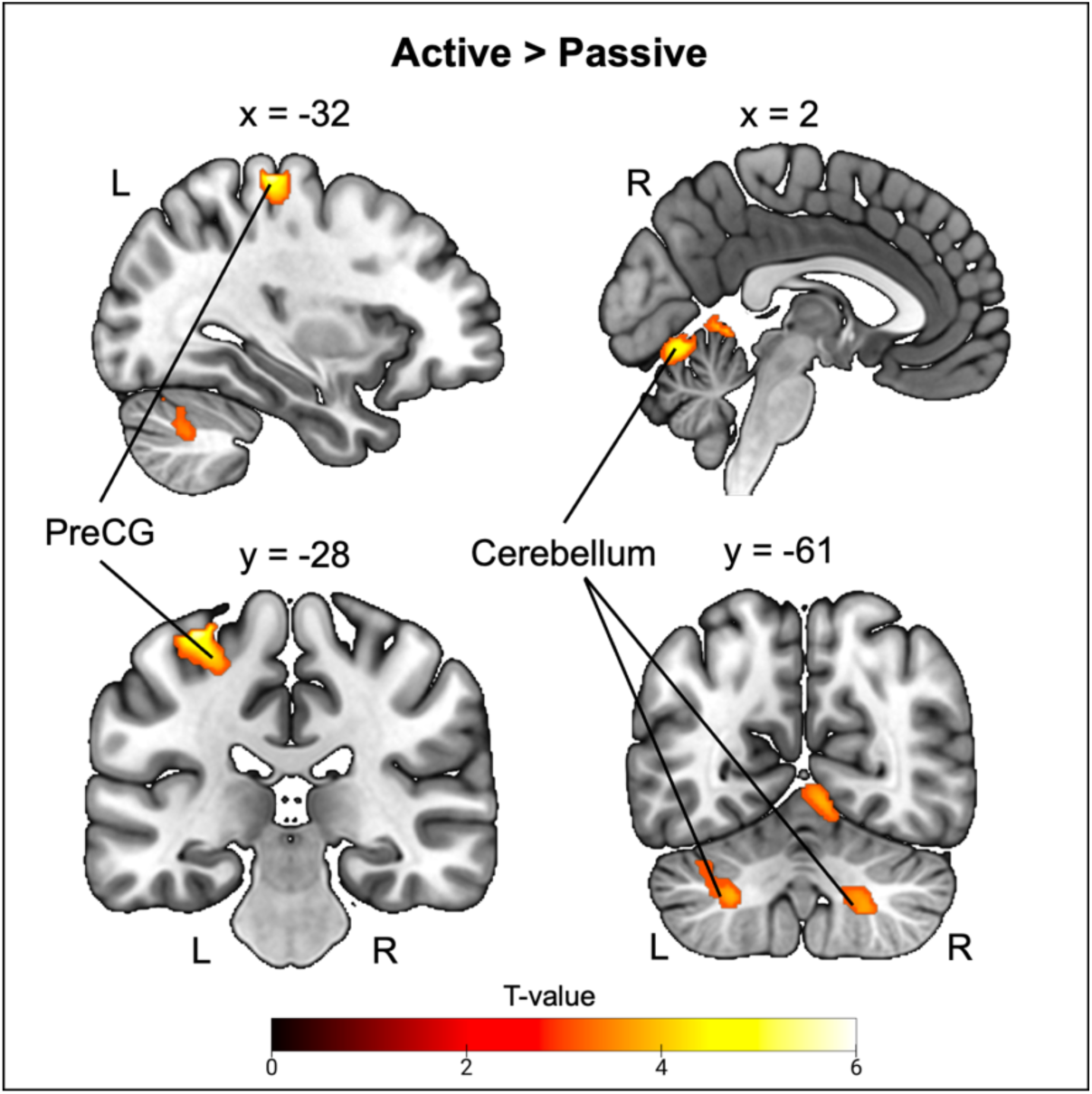
Group results for the main effect of movement type (Active > Passive). In test phases, active movements were associated with increased activity in left precentral gyrus and bilateral cerebellum. PreCG = precentral gyrus, L = left, R = right.

**Supplementary Table 4.**
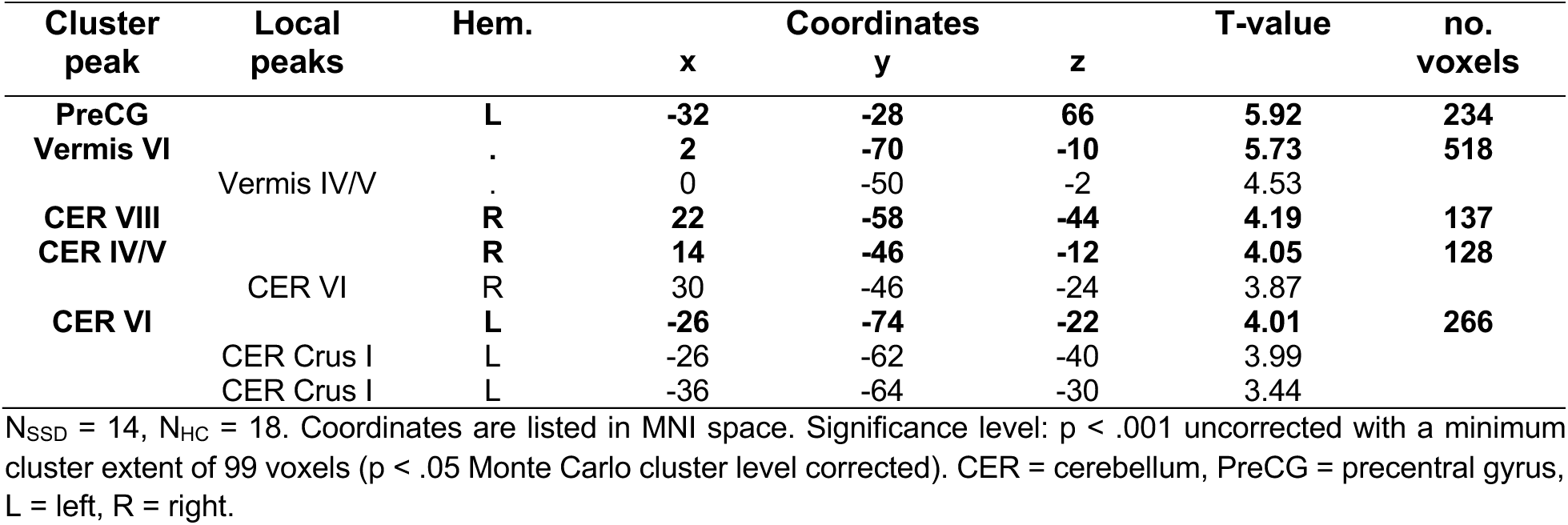
Group results for the main effect of movement type (Active > Passive).

### S5. Details on correlation analyses between recalibration effects and symptom severity

Exploratory correlation analyses were performed to investigate the association between symptom severity, assessed via the SAPS total score and subscores for hallucinations and delusions, and both behavioral and neural TREs. A summary of all correlation results is provided in **Supplementary Table 5**. Scatter plots for significant correlations are provided in **Supplementary Fig. 2**.

**Supplementary Table 5.**
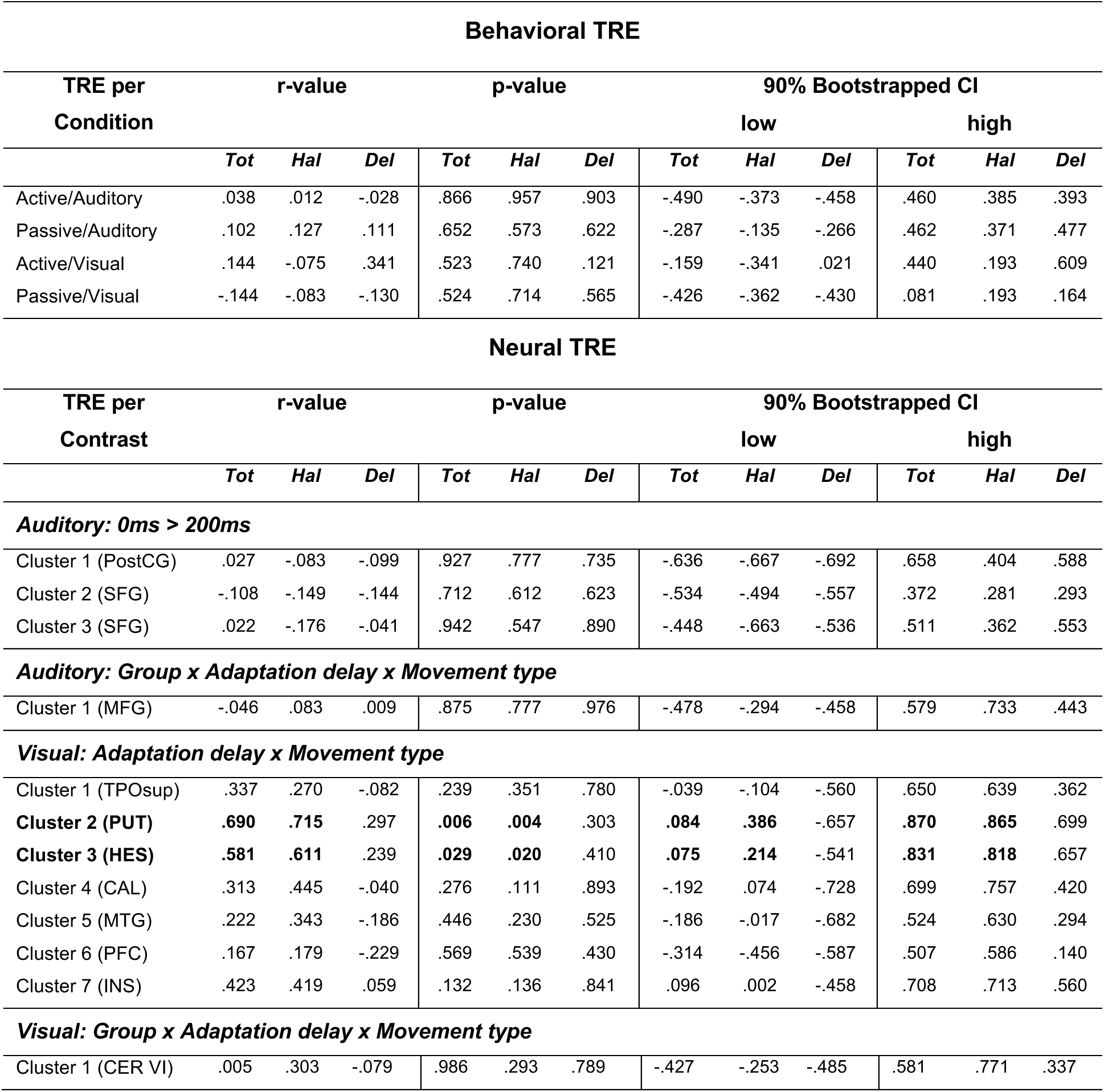
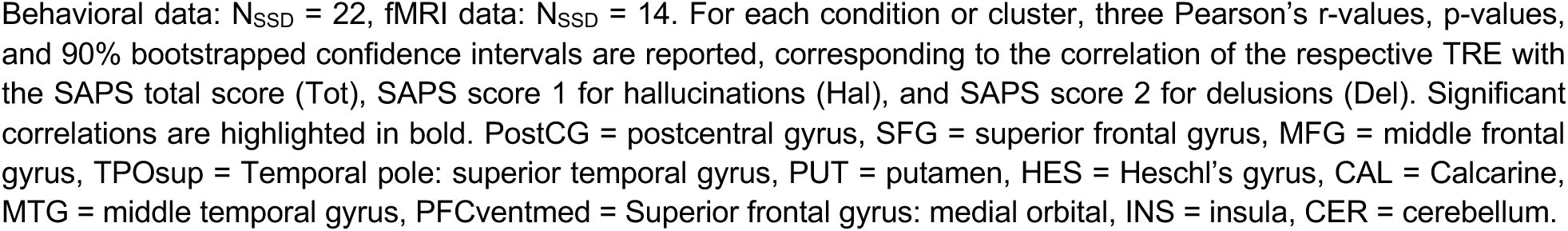
Correlations between behavioral and neural TREs and symptom severity.

**Supplementary Fig. 2.**
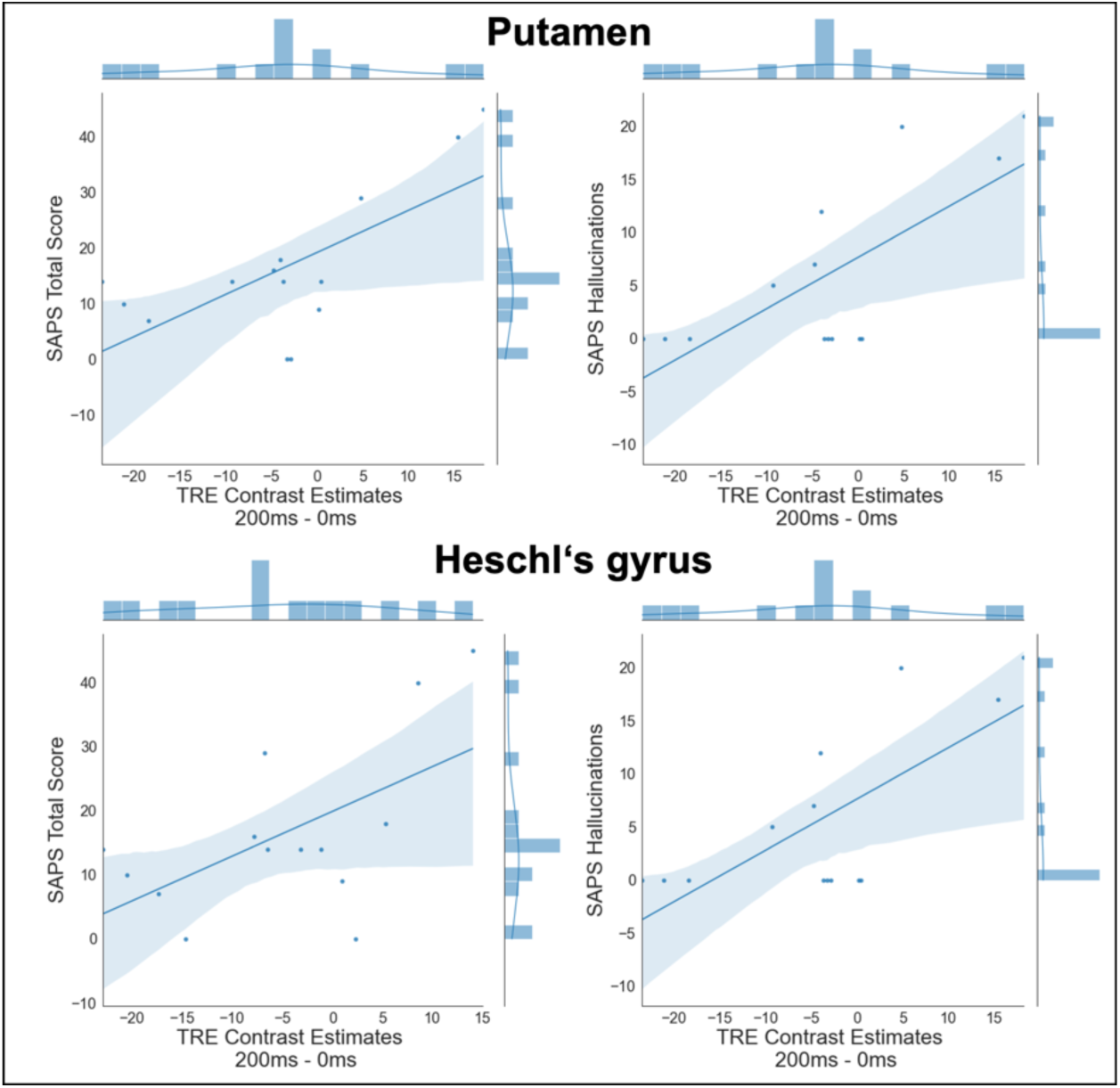
Scatter plots for significant correlations of TREs and symptom severity. Patients with a higher SAPS total score and a higher hallucinations subscore were the ones that deviated most strongly from the overall activation pattern in active conditions (i.e., a negative neural TRE) observed across both groups in putamen and Heschl’s gyrus (visual test modality; contrast: Adaptation delay x Movement type).

### S6. Exploratory analysis of delay-dependent activations during adaptation phases

We used a parametric analysis to identify brain regions which experienced a linear change in activation throughout the adaptation phases. This analysis relied on the same preprocessed data that were used in the main manuscript and on the same definition of regressors in the GLM for the experimental conditions and events of no interest. As described in the main manuscript, for the adaptation phases, regressors were defined based on eight experimental conditions comprising the factors *adaptation delay* (0ms, 200ms), *movement type* (active, passive), and *adaptation phase* (early, late). To parametrically modulate these regressors, the number of button presses performed during early adaptation phases (i.e., before the presentation of the fixation cross) and during late phases (i.e., after the fixation cross) were used as parametric regressor. For the single-participant GLMs, T-maps were generated by contrasting each of the eight parametric regressors against the implicit baseline. For group-level analyses, the resulting contrast estimates from each participant were entered into a flexible factorial design. As in the analysis reported in the main manuscript, we were interested in main and interaction contrasts involving the *adaptation delay* factor.

We first examined differences in parametric brain activation changes across both groups. Here, the significant interaction contrast composed of the factors *adaptation delay* and *movement type* revealed that activation in right supplementary motor area, right caudate, and in left cerebellum decreased linearly throughout adaptation when participants were exposed to the 200ms delay (see **Supplementary Fig. 3** and **Supplementary Table 6**). This effect was comparatively stronger in active movement conditions than in passive ones. The SMA is thought to be involved in generating the efference copy of motor commands, which is then projected to the cerebellum. The cerebellum uses this efference copy to perform forward model operations by predicting the sensory outcomes of the motor commands. If necessary, the prediction error is then sent back to the SMA to adjust the motor commands for future actions.^10,11^ It has already been shown before that the connectivity between the cerebellum and the SMA temporarily increases when we are exposed to an additional delay between action and action-outcome.^12^ Therefore, it is likely that at the beginning of the adaptation phases, a stronger prediction error signal is present in the cerebellum and SMA, which then gradually diminishes as the forward model integrates the constant delay into its predictions.

To investigate whether parametric brain activation changes during adaptation differed between patients with SSD and HC, we examined contrasts involving the factors *group* and *adaptation delay*. A significant four-way interaction of *group*, *adaptation delay*, *movement type* and *adaptation phase* emerged, with a cluster spanning left precuneus and right posterior cingulate cortex. In these regions, HC showed a linear increase of activation in late adaptation phases while exposed to the 200ms delay, with a stronger effect in active than in passive conditions. For SSD, a similar activation increases in these regions could be observed during early adaptation phases. Both regions are part of a network of cortical midline structures which are involved in processing self-referential information and have been associated with the experience of agency.^13,14^ It appears that these regions show stronger activation in HC when they perceive synchrony between action and outcome or when the outcome is attributed to their own action, which is likely to occur in later phases of adaptation.^15^ In SSD, functional alterations in the precuneus during self-referential processing are well-documented^16^ and may, in this case, lead to deviations in the process of attributing the delayed outcome to the own action compared to HC. However, the precise nature of these activity deviations and their potential consequences remain open based on the present data.

**Supplementary Fig. 3.**
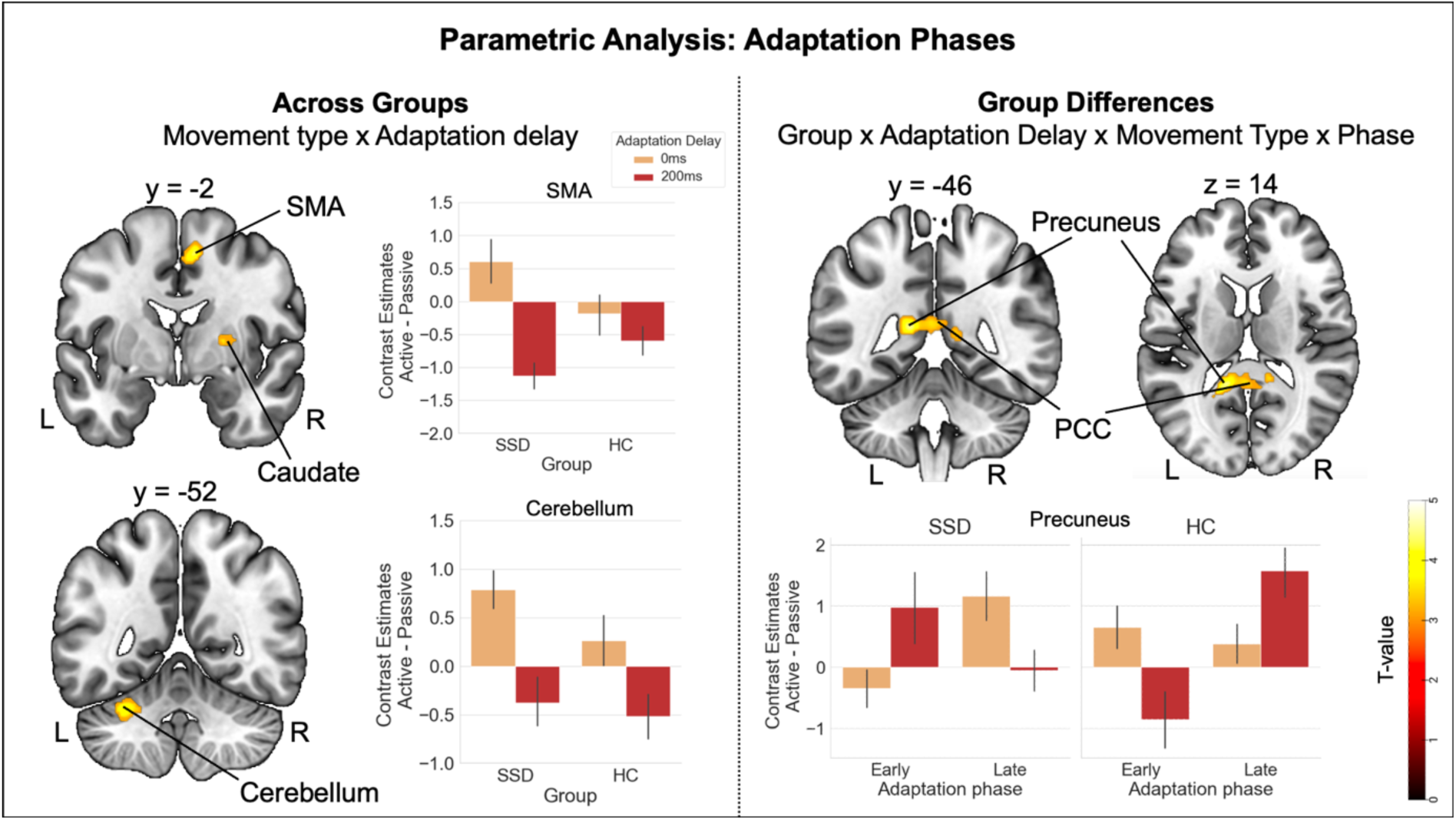
Group results for the parametric analysis of adaptation phases. Left panel: Parametric analyses across groups showed a decrease in activation over the course of adaptation to the 200ms delay in right supplementary motor area, right caudate, and left cerebellum. This effect was comparatively stronger in active movement conditions than in passive ones. **Right panel:** Group differences emerged in left precuneus and right posterior cingulate cortex where HC show an activation increase in late adaptation phases while exposed to the 200ms delay, with a stronger effect in active than in passive conditions. For SSD, this activation increase occurred during early adaptation phases. For visualization, contrast estimates (eigenvariates extracted with the VOI function of SPM) are displayed as the difference between active and passive conditions, emphasizing the extent to which the effect is more pronounced in active compared to passive conditions. Error bars show standard errors of the mean. SMA = supplementary motor area, PCC = posterior cingulate cortex, L = left, R = right.

**Supplementary Table 6.** Group results for the parametric analysis of adaptation phases.

| Cluster peak | Local peaks | Hem. | x | Coordinates<br>y | z | T-value | no. voxels |
| --- | --- | --- | --- | --- | --- | --- | --- |
| **Across Groups: Interaction Adaptation delay x Movement type** |  |  |  |  |  |  |  |
| **SMA** |  | **R** | **8** | **-2** | **54** | **4.42** | **103** |
| **CAU** |  | **R** | **18** | **8** | **16** | **4.22** | **132** |
|  | PUT | R | 24 | 0 | 6 | 3.83 |  |
| **CER VI** |  | **L** | **-30** | **-52** | **-28** | **4.19** | **100** |
|  | CER IV/V | L | -18 | -48 | -30 | 3.35 |  |
| **Interaction Group x Adaptation delay x Movement type x Phase** |  |  |  |  |  |  |  |
| **PCUN** |  | **L** | **-14** | **-46** | **14** | **4.60** | **342** |
|  | PCC | R | 12 | -42 | 10 | 3.85 |  |
|  | PCC | R | 4 | -36 | 6 | 3.75 |  |

N<sub>SSD</sub> = 14, N<sub>HC</sub> = 18. Coordinates are listed in MNI space. Significance level: $p < .001$ uncorrected with a minimum cluster extent of 99 voxels ( $p < .05$ Monte Carlo cluster level corrected). SMA = supplementary motor area, CAU = caudate, PUT = putamen, CER = cerebellum, PCUN = precuneus, PCC = posterior cingulate cortex, L = left, R = right.

### S7. Exploratory correlation analyses between chlorpromazine (CPZ) equivalents and behavioral and neural recalibration effects

To investigate whether the observed effects were influenced by antipsychotic medication, we conducted exploratory correlations between the CPZ equivalents (mg/d) and both behavioral and neural TREs. To convert the medication doses into CPZ equivalents, we used the “chlorpromazineR” package (version 0.2.0) for R^17^ (version 4.2.2). The conversion factors of aripiprazole, quetiapine, olanzapine, clozapine, amisulpride, risperidone, paliperidone, and flupenthixol were based on the work of Gardner et al. (2010)^18^, while the conversion factors for melperone and cariprazine were taken from Leucht et al. (2016)^19^ and Leucht et al. (2020),^20^ respectively. On the behavioral level, CPZ equivalents (*Mean* = 419.294, *SD* = 435.893) were then correlated with the TREs for each experimental condition. On the neural level, CPZ equivalents (*Mean* = 426.633, *SD* = 518.425) were correlated with the TREs of the contrast estimates (200ms – 0ms) of each cluster for significant fMRI contrasts involving the *adaptation delay* factor following a similar logic as described for the correlation analyses with symptom severity reported in the main manuscript. None of the correlations reached significance (see **Supplementary Table 6** for a detailed summary of all correlation results), suggesting that the patients’ medication did not interfere with recalibration processes and performance in the task.

**Supplementary Table 6.** Correlations between behavioral and neural TREs and CPZ equivalents.

| **Behavioral TRE** |  |  |  |  |
| --- | --- | --- | --- | --- |
| **TRE per Condition** | **r-value** | **p-value** | **90% Bootstrapped CI** |  |
|  |  |  | **low** | **high** |
| Active/Auditory | .135 | .550 | -.110 | .403 |
| Passive/Auditory | -.097 | .668 | -.468 | .156 |
| Active/Visual | .089 | .693 | -.184 | .326 |
| Passive/Visual | .208 | .354 | -.202 | .616 |
| **Neural TRE** |  |  |  |  |
| **TRE per Contrast** | **r-value** | **p-value** | **90% Bootstrapped CI** |  |
|  |  |  | **low** | **high** |
| ***Auditory: 0ms > 200ms*** |  |  |  |  |
| Cluster 1 (PostCG) | .304 | .290 | -.025 | .542 |
| Cluster 2 (SFG) | .498 | .070 | .081 | .779 |
| Cluster 3 (SFG) | .240 | .409 | -.293 | .645 |
| ***Auditory: Group x Adaptation delay x Movement type*** |  |  |  |  |
| Cluster 1 (MFG) | .257 | .376 | -.509 | .861 |
| ***Visual: Adaptation delay x Movement type*** |  |  |  |  |
| Cluster 1 (TPOsup) | .011 | .971 | -.433 | .482 |
| Cluster 2 (PUT) | -.272 | .347 | -.746 | .232 |
| Cluster 3 (HES) | -.254 | .380 | -.662 | .306 |
| Cluster 4 (CAL) | -.193 | .509 | -.771 | .423 |
| Cluster 5 (MTG) | -.075 | .798 | -.730 | .481 |
| Cluster 6 (PFC) | -.079 | .789 | -.595 | .500 |
| Cluster 7 (INS) | -.337 | .239 | -.788 | .144 |
| ***Visual: Group x Adaptation delay x Movement type*** |  |  |  |  |
| Cluster 1 (CER VI) | -.234 | .421 | -.703 | .757 |

Behavioral data: $N_{SSD} = 22$ , fMRI data: $N_{SSD} = 14$ . For each condition or cluster, Pearson's r-values, p-values, and 90% bootstrapped confidence intervals are reported, corresponding to the correlation of the respective TRE with the patients' CPZ equivalents. PostCG = postcentral gyrus, SFG = superior frontal gyrus, MFG = middle frontal gyrus, TPOsup = Temporal pole: superior temporal gyrus, PUT = putamen, HES = Heschl's gyrus, CAL = Calcarine, MTG = middle temporal gyrus, PFCventmed = Superior frontal gyrus: medial orbital, INS = insula, CER = cerebellum.

